# Bypass of an epigenetic S-phase transcriptional module by convergent nutrient stress signals

**DOI:** 10.1101/741363

**Authors:** Marie Delaby, Gaël Panis, Coralie Fumeaux, Laurence Degeorges, Patrick H. Viollier

## Abstract

The signals feeding into bacterial S-phase transcription are poorly understood. Cellular cycling in the alpha-proteobacterium *Caulobacter crescentus* is driven by a complex circuit of at least three transcriptional modules that direct sequential promoter firing during the G1, early and late S cell cycle phases. In alpha-proteobacteria, the transcriptional regulator GcrA and the CcrM methyltransferase epigenetically activate promoters of cell division and polarity genes that fire in S-phase. By evolving *Caulobacter crescentus* cells to cycle and differentiate in the absence of the GcrA/CcrM module, we discovered that phosphate deprivation and (p)ppGpp alarmone stress signals converge on S-phase transcriptional activation. The cell cycle oscillations of the CtrA protein, the transcriptional regulator that implements G1 and late S-phase transcription, are essential in our evolved mutants, but not in wild-type cells, showing that the periodicity in CtrA abundance alone can sustain cellular cycling without GcrA/CcrM. While similar nutritional sensing occurs in other alpha-proteobacteria, GcrA and CcrM are not encoded in the reduced genomes of obligate intracellular relatives. We thus propose that the nutritional stress response induced during intracellular growth obviated the need for an S-phase transcriptional regulator.

## Introduction

The alpha-proteobacterial lineage offers a tractable system to elucidate which regulatory genes are necessary to sustain cellular cycling in a primordial (bacterial) cell. *Caulobacter crescentus* like many alpha-proteobacteria undergoes an asymmetric cell division^1^, giving two progeny cells with distinct morphologies and fates: the capsulated stalked cell that resides in S-phase and the motile and piliated swarmer cell that resides in a G1-like non-replicative state and must differentiate into a St cell before proceeding to division (Fig. 1A). Synchronization of these cell types is possible on the basis of a difference in buoyancy conferred by the cell cycle regulated capsule which allows the enrichment of G1 cells. During the G1 to S cell transition, the polar pili and chemosensory machines are lost^2^, the flagellum is shed and a stalk elaborates from the vacated old cell pole. Coincident with the morphological changes, DNA replication competence is acquired, and chromosome replication commences bidirectionally from the single origin of replication (C*ori*). During S-phase, the cell matures into an asymmetric pre-divisional cell decorated with a new flagellum, chemosensory and pilus secretion complexes assembled at the pole opposite the stalk. Once chromosome replication and segregation is completed, cytokinesis partitions the two daughter cells and triggers the activation of dissimilar cell fate transcriptional programs in each progeny^3^ (Fig. 1B).

**Figure 1.**
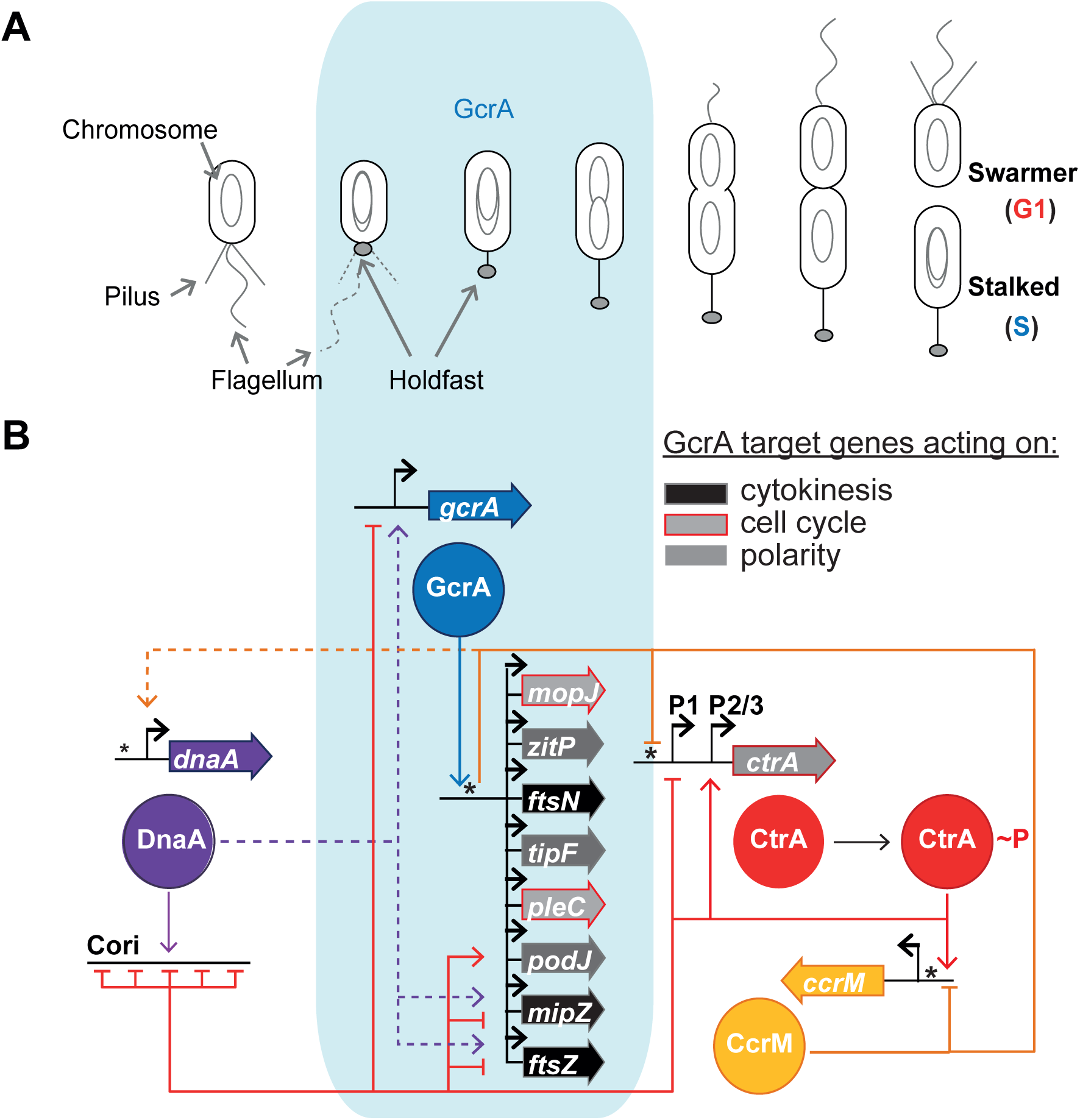
Model of *C. crescentus* cell cycle and S-phase regulatory interactions controlled by the GcrA/CcrM module. (**A**) Schematic of *C. crescentus* cell cycle showing the chromosome replication cycle and polar remodeling. The blue box delimits the early S-phase period (theta structure of chromosome) controlled by the GcrA transcriptional factor. (**B**) Schematic of the regulatory interactions controlling cycle-regulated promoters. In early S-phase, the GcrA/CcrM module activates promoters of genes implicated in cytokinesis (boxed in black), polarity (dark grey) and cell cycle regulation (light grey). Blue line represents the binding of GcrA to early S-phase promoters. Orange lines represent methylation control by CcrM on GcrA target promoters with the 5′-GANTC-3′ methylation motif (indicated as asterisk). Red lines represent direct transcriptional control by CtrA (activation or repression shown as arrows or T-bar, respectively). Possible control of DnaA at the promoters of *gcrA*, *podJ* and *ftsZ* is depicted as dotted lines (purple). Note that the DnaA occupancy at these promoters has only be demonstrated *in vitro*, but not yet been observed *in vivo*.

Cell cycle progression (chromosomal replication and cytokinesis) is coordinated with differentiation via a complex, cyclical regulatory circuit in which global transcriptional regulators sequentially activate promoters. The G1-phase transcriptional program is controlled by the conserved and essential master cell cycle transcriptional regulator CtrA^3^, a DNA-binding response regulator that usually activates promoters that fire in G1-phase or in late S-phase. By contrast, the transcriptional regulator GcrA is only present in early S-phase and binds promoters of genes encoding cytokinetic, polarity or regulatory proteins such as CtrA required for subsequent cell cycle stages (Fig. 1B).

GcrA is a conserved but non-canonical transcriptional regulator that interacts with the RNA polymerase (RNAP) sigma 70 (σ^70^) via domain 2 that binds to the -10 promoter elements^4, 5^. Unlike canonical transcription factors that bind to promoters and then recruit RNAP holoenzyme, epitope-tagged GcrA was proposed to first bind RNAP through its interaction with σ^70^ and is then placed at σ^70-^dependent promoters by σ^70^•RNAP ^4^. An untested corollary of this model is that GcrA depends on σ^70^•RNAP for proper positioning at σ^70^ target promoters, rather than σ^70^•RNAP recruitment at target promoters depending on GcrA.

Paradoxically, earlier ChIP-Seq experiments of (unmodified) GcrA precipitated from wild-type cells using polyclonal antibodies to GcrA had revealed preferential binding of GcrA to a subclass of σ^70^-dependent promoters^6, 7^. Since binding of GcrA to such a target promoter had been recapitulated *in vitro* without σ^70^ and RNAP, GcrA can bind DNA specifically. The basis for the promoter selectivity was shown to reside in the epigenetic (methylation) mark in the context of the 5’-GANTC-3’ sequence found in many, but not all, σ^70^-dependent promoters. The adenine (N6) methylation mark is introduced by the CcrM adenine methyltransferase^7, 8^. Preferential promoter binding of GcrA is no longer observed^6, 7^ in the absence of CcrM. Since preferential binding of methylated target promoters by GcrA is not commensurate with general σ^70^•RNAP promoter associations, the σ^70^•RNAP interaction likely serves to recruit GcrA to σ^70^-dependent promoters. This would allow a firm association of GcrA at promoters having a CcrM-dependent methylation mark to stabilize σ^70^•RNAP at the promoter and induce promoter firing.

Loss of GcrA and/or CcrM lead to similar severe pleiotropic defects, rendering cells non-motile, unsynchronizable (because of a defect in capsule control), strongly elongated with multiple chromosomes^7, 9^. Selection of efficiently dividing Δ*gcrA* cells by transposon (Tn) mutagenesis had yielded promoter-up mutations in the GcrA-target promoter of the *ftsN* cell division gene conferred by an outward directed promoter within the *himar1* Tn (P*_ftsN_*::Tn). Although P*_ftsN_*::Tn directs FtsN expression independently of GcrA^7^ and ameliorates division, other defects of Δ*gcrA* P*_ftsN_*::Tn cells are not corrected, indicating that cellular cycling is not restored. Thus, the question whether a proper *C. crescentus* cell cycle can be sustained without GcrA (and CcrM) remains unresolved. The GcrA/CcrM module is co-conserved in the alpha-proteobacteria, with the notable exception of the obligate intracellular *rickettsial* lineage, suggesting that at least rickettsial cells can cycle in the absence of GcrA and CcrM.

Here we show that a combination of phosphate starvation and [(p)ppGpp] alarmone stress signalling allows *C. crescentus* cells to cycle without GcrA (and CcrM). We further show that under these conditions S-phase promoters can fire independently of GcrA and CcrM. Moreover, orthologous S-phase promoters from *Sinorhizobium meliloti* and members of the obligate intracellular *Rickettsiales* (*Rickettsia prowazekii* and *Ehrlichia canis*) also answer to phosphate starvation, either in *C. crescentus* or in *S. meliloti*. Using ChIP experiments we found that σ^70^ occupancy is reduced at GcrA target promoters in Δ*gcrA* cells, supporting a model in which GcrA binds and stabilizes σ^70^-containing RNAP holoenzyme at methylated S-phase promoters, inducing efficient firing in the absence of nutrient stress.

## Results

### Robust and asymmetric cellular cycling without GcrA

We evolved cells to cycle and differentiate efficiently in the absence of GcrA by extended incubation of Δ*gcrA*::Ω Δ*gcrB* parental cells on swarm (0.3%) agar plates. In the absence of GcrA, cells manifest a defect in the production of properly differentiated (motile) G1-phase cells (Fig. 2A and 2C) due to an insufficiency in expression of G1-phase regulator CtrA, several polarity determinants (PodJ, ZitP, TipF) and cell division proteins (FtsN, MipZ; Fig. 1B, 2D and S1B), all of which are expressed from a GcrA-activated promoter in early S-phase (Fig. 1B). As some cell cycle mutants exhibit a swarming defect owing to a reduction of motile G1 cells, we reasoned that restoration of the cell cycle defect and production of G1 cells can be obtained by evolving Δ*gcrA* mutant cells on swarm agar. Since *C. crescentus* encodes an ortholog of GcrA (GcrB), we used Δ*gcrA*::Ω Δ*gcrB* parental cells to avoid isolating gain-of-function mutations in the *gcrB* gene that might otherwise compensate for the absence of GcrA. After 6 days of incubation of Δ*gcrA*::Ω Δ*gcrB* on swarm agar at 30°C, we isolated three independent mutant clones from motile flares (Fig. 2A). Phase contrast microscopy of the three motile suppressor mutant cultures revealed short (swimming) cells, rather than the long non-motile cells typically seen in Δ*gcrA*::Ω Δ*gcrB* parental cultures (Fig. 2B). Importantly, we observed a recovery of a substantial G1 cell population as determined in flow cytometry experiments using Fluorescence-activated cell sorting (FACS, Fig. 2C, S1A). At the molecular level, all three suppressor mutant clones express CtrA, PodJ, ZitP, TipF, MipZ and FtsN to near *WT* levels or beyond (Fig. 2D, S1B) and were able to export again the FljK flagellins (Fig. 2D), indicating that they are assembled into the extracellular flagellar filament (that can be separated from cells mechanically by shearing). We also observed that, unlike Δ*gcrA*::Ω Δ*gcrB* parental cells, the three motile suppressors mutants can again implement the cell cycle-regulated switch in capsulation (scored as change in buoyancy, Fig. S1C), promote cycling of the cell cycle regulated proteins such as CtrA and PodJ (Fig. S2A) in concert with the progression of the chromosome replication (Fig. S2B) and can again be infected by the S-layer-specific phage φCr30^10^ and the pilus-specific phage φCbK^11^ (Fig. S1C). Expression of the S-layer subunit protein RsaA^12^ was previously reported to be impaired upon GcrA depletion^4, 7^ and pilus assembly is dependent on the ZitP and PodJ polarity proteins that are expressed from GcrA-dependent S-phase promoters^7, 13^ (Fig. 1B). Interestingly, despite these improvements in cell cycle control and differentiation (Fig. S2A, S2B), the doubling time (T_d_) of the three motile suppressor mutants is increased compared to that of the Δ*gcrA*::Ω Δ*gcrB* parent (Fig. 2B, S1D), pointing to a phenotypic trade-off arising from the forced selection of cell cycle control in the absence of GcrA/B. In sum, our findings show that despite the surge in expression of cell cycle genes in early S-phase by GcrA, it is possible for *Caulobacter* cells to cycle robustly and asymmetrically in the absence of GcrA/B.

**Figure 2:**
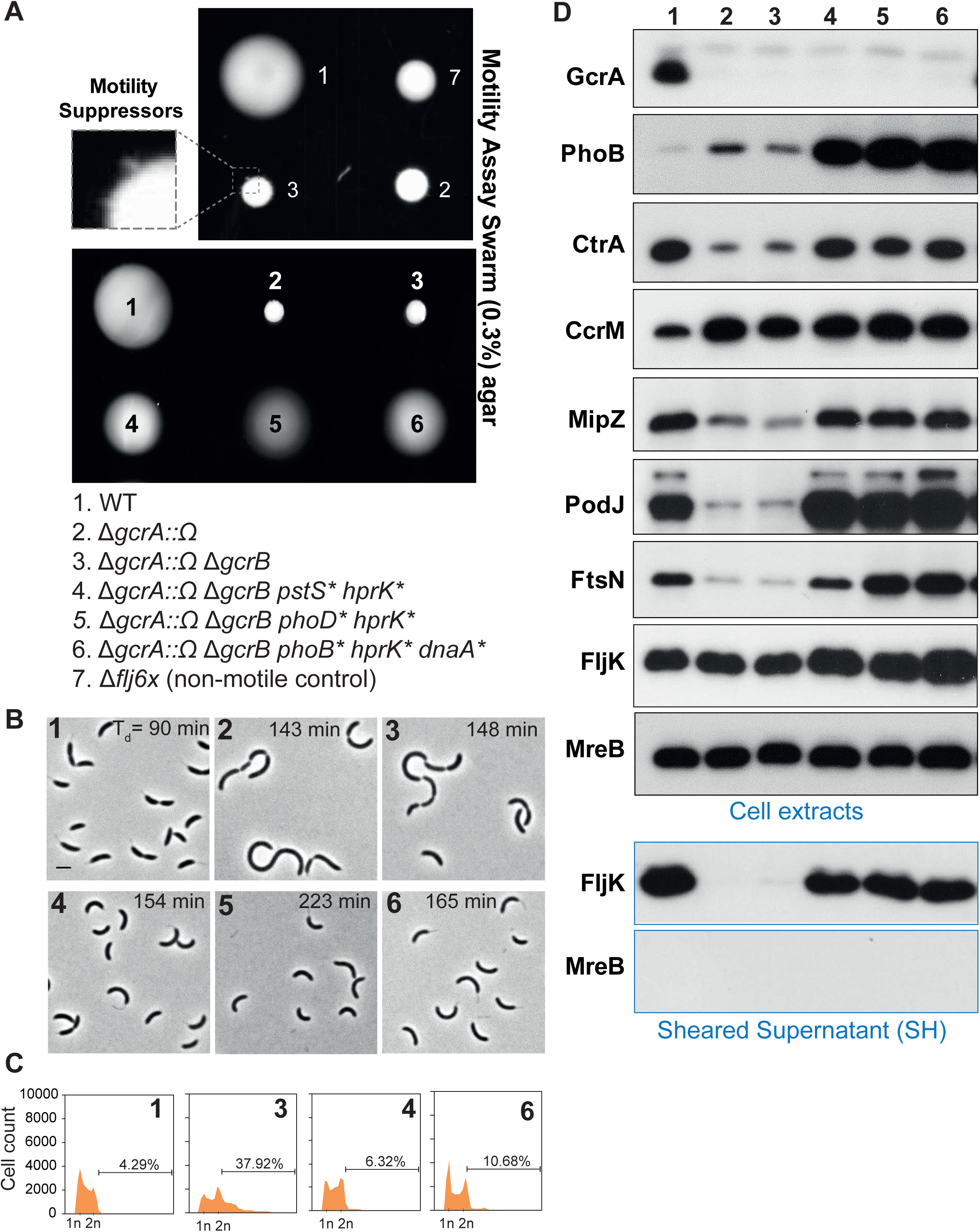
Isolation and characterization of Δ*gcrA* motility suppressors. (**A**) Motility assays on swarm (0.3 %) agar of *WT*, Δ*fljx6,* Δ*gcrA::*Ω and Δ*gcrA::*Ω Δ*gcrB*. The square points to the emergence of spontaneous motile suppressors from the Δ*gcrA::*Ω Δ*gcrB* double mutant. Three spontaneous Δ*gcrA::*Ω Δ*gcrB* motility suppressors were isolated and their genomes sequenced, revealing the following point mutations: (i) in *hprK^D47E^* and (ii) in the phosphate uptake pathway (either *pstS^G61S^*(4), *phoD^V117A^*(5) or *phoB^T74A^* (6)). The last suppressor (6) had a third mutation in *dnaA*^H334R^. (**B**) Phase microscopy of cells grown exponentially in PYE. Values indicate the generation doubling time (from three biological replicates) and scale bar represents 2 μm. (**C**) DNA content analysis as determined by FACS (FL1-A channel) of *WT* and Δ*gcrA::*Ω mutant cells and derivatives grown in PYE to exponential phase. Percentages of cells containing >2 chromosomes are also indicated. (**D**) Immunoblots showing steady-state levels of indicated proteins in extracts of *WT* and Δ*gcrA::*Ω mutant cells grown in PYE to exponential phase. FljK abundance released into culture supernatants after shearing, as a marker for flagellation, is also indicated. Cytoplasmic MreB actin serves as a negative loading control for possible cell lysis that could have occurred during shearing of the cells.

### Phosphoenolpyruvate phosphotransfer cues into the cell cycle

Whole genome sequencing of the three motile Δ*gcrA*::Ω Δ*gcrB* suppressors revealed several mutations, including a common missense mutation within *hprK* (*CCNA_00239*) that is not present in the Δ*gcrA*::Ω Δ*gcrB* parental strain. This mutation (*hprK^D47E^*) changes the conserved aspartic acid at position 47 into a glutamic acid (D47E) in the serine kinase/phosphatase HprK, a key regulator of the phosphoenolpyruvate phosphotransferase system (PTS, Fig. 3A). In alpha-proteobacteria, the PTS mediates the transfer of a phosphoryl group from phosphoenolpyruvate to stimulate the accumulation of the alarmone guanosine (penta) tetra-phosphate commonly referred to as (p)ppGpp. The phosphoryltransfer proceeds via EI^Ntr^/PtsP→HPr→EII culminating in the regulation of the bifunctional (p)ppGpp synthase/hydrolase SpoT^14, 15^ (Fig. 3A). HprK is thought to phosphorylate serine 49 residue of Hpr, thereby preventing phosphorylation of Hpr (∼P) at histidine 18 (H18) by the EI^Ntr^/PtsP protein and thus to interfere with the EII-dependent stimulation of SpoT^16^. Recently, both mutations in the PTS EI^Ntr^/PtsP and HprK proteins were also identified during *in vitro* evolution experiments in PYE broth selecting for mutations that improve the growth rate of Δ*ccrM* cells^9^. It is important to note that in the Δ*ccrM* evolution experiments a strong selection was imposed for rapid growth/division, rather than proper cellular cycling as our selection with swarm agar. Consistent with this shorter doubling time, the *hprK* (*hprK1.4*) mutation surfacing in Δ*ccrM* cells also promoted an increase in transcription of the *mipZ* and *ftsZ* cell division genes whose promoters are known to be GcrA-dependent. In support of this, transposon insertion sequencing (Tn-Seq) analysis of Δ*ccrM*::Ω cells mutagenized with a *himar1* transposon (Tn) implicated the PTS system including the *hprK* gene (ranked 22^nd^) and *CCNA_00240* (ranked 59^th^, PTS system IIA component) in regulating growth of Δ*ccrM*::Ω cells compare to *WT* (Fig. S3A, S3B, Table S1). Moreover, the *spoT* gene (ranked 14^th^) encoding the unique bifunctional (p)ppGpp synthase/hydrolase was also overrepresented as a Tn insertion promoting growth of Δ*ccrM*::Ω cells. Importantly, the Tn insertions occurred outside of the synthase domain. Finally, Tn insertions in the region upstream of *ftsZ* were also found to be greatly overrepresented in Δ*ccrM*::Ω cells compared to the WT (Fig. S3C, Table S1). In a comparable experiment where we previously selected for rapid (colonial) growth after transposon mutagenesis of Δ*gcrA*::Ω Δ*gcrB* cells, Tn insertions causing an upregulation in expression of the *ftsN* cell division gene, whose promoter is also a GcrA target, were isolated. However, both these selections for improved growth differ from our selection for efficient cell cycle (motility) control imposed on Δ*gcrA*::Ω Δ*gcrB* cells in swarm agar from which is suited to evolve near normal cellular cycling properties.

**Figure 3.**
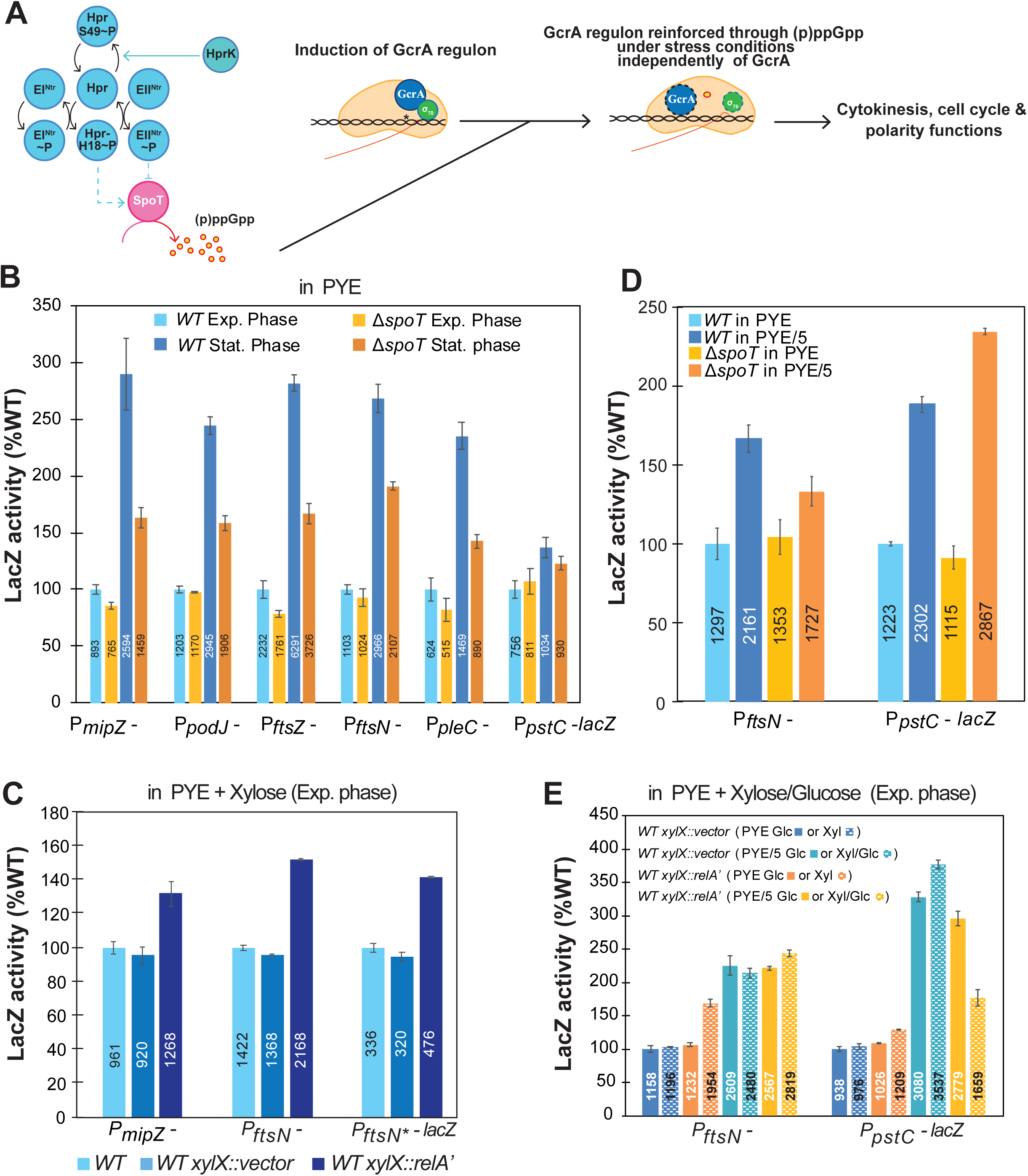
The PTS system, the (p)ppGpp alarmone and phosphate starvation affect the transcription of genes also controlled by GcrA. (**A**) Schematic model of the network connecting the phosphoenolpyruvate phosphotransferase system (PTS) system, the (p)ppGpp alarmone and the transcription of GcrA targeted genes. EI^Ntr^ corresponds also to PtsP protein. (**B**) β-galactosidase produced from various promoter-probe plasmids in which either the *mipZ*, *podJ*, *ftsZ*, *ftsN, pleC* or *pstC* promoter is fused to a promoterless *lacZ* gene was measured in *WT* and the Δ*spoT* exponential (exp.) or stationary (stat.) phase cells grown in PYE. (**C**) β-galactosidase activities of P*_mipZ_-*, P*_ftsN_-* and P*_ftsN*_-lacZ* fusions, in *WT* cells carrying or not *relA’* under control of the xylose-inducible P*_xylX_*-promoter integrated at the *xylX* locus, were measured after 5 hours of growth in PYE supplemented with 0.3% of Xylose. (**D**) β-galactosidase activities of P*_ftsN_-* or P*_pstC_-lacZ* fusions in *WT* and Δ*spoT* strains cultivated to exponential phase in PYE or in phosphate-limited PYE/5. The P*_pstC_*-*lacZ* promoter reporter serves a positive control for the phosphate-limited conditions. (**E**) β-galactosidase activities of P*_ftsN_-* and P*_pstC_-lacZ* fusions in *WT* carrying or not *relA’* under control of the xylose-inducible P*_xylX_*-promoter integrated at the *xylX* locus were measured after 5 hours of growth in PYE or PYE/5 supplemented with 0.3% of Xylose or/and 0.2% of Glucose (to repress P*_xylX_*) as indicated. (**B**, **C**, **D** and **E**) β-galactosidase activities expressed in percentage relative to *WT* measured as Miller units. Data are from four independent experiments; error bars are standard deviation.

Since HprK is an indirect regulator of SpoT and since we found the *hprK^D47E^* mutation in all three independent motile suppressor mutants, we wondered if GcrA-dependent promoters in general are responsive to elevated (p)ppGpp levels. This possibility seemed likely because the levels of GcrA-dependent proteins (including MipZ, ZitP, PodJ and FtsN) examined by immunoblotting were elevated relative to the Δ*gcrA*::Ω Δ*gcrB* parent (Fig. 2D, S1B) and because we recently reported that the GcrA-dependent *tipF*, *ctrA* and *mopJ* promoters are induced is stationary phase in a SpoT-dependent manner^17^. Indeed, *lacZ*-based promoter probe assays (Fig. 3B) showed that all GcrA-dependent promoters examined exhibit a dependency on (p)ppGpp synthesized by SpoT in stationary phase. Moreover, we found that ectopic production of (p)ppGpp in exponential phase (triggered by transcriptional induction of the heterologous (p)ppGpp-synthase RelA’ from *Escherichia coli*) leads to an increase in β-galactosidase activity in *C. crescentus WT* cells harbouring *lacZ*-based promoter probe reporters of GcrA (P*_ftsN_*-*lacZ* and P*_mipZ_*-*lacZ*, Fig. 3C). Since we found that P*_ftsN_*\*-*lacZ* reporter (in which the single GANTC methylation motif in the promoter is mutated) still responds to an increase in (p)ppGpp levels (Fig. 3C), we conclude that the suppressive mechanism of promoter control operates independently of GcrA/B, as already suggested by the motility suppressor selection described above. Moreover, the induction of P*_ftsN_*\*-*lacZ* in medium that has been diluted five-fold (PYE/5) with water containing 0,3% glucose was lost in Δ*spoT* cells (Fig. S4A), showing that induction of the GcrA-dependent promoters by (p)ppGpp can occur without a methylated enhancer sequence in the GcrA target promoter.

The finding that all three suppressor mutants exhibit a high level of expression of many GcrA-dependent genes, prompted us to test if reintroducing GcrA and thereby further augmenting expressing early S-phase gene expression would be beneficial or harmful to our suppressor cells. We found that a GcrA-expression plasmid (in which GcrA is expressed from vanillate-inducible P*_van_* promoter on the pMT335 plasmid, pMT335-*gcrA*) did not have a beneficial affect and was in fact less well tolerated by the three suppressor mutants, even in the absence of inducer (vanillate) as determined by efficiency of plating experiments (Fig. S5A and S5B). By contrast, pMT335-*gcrA* restored plating efficiency to Δ*gcrA*::Ω Δ*gcrB* cells and did not adversely affect that of *WT.* We interpret this defect as a consequence of further induction of the GcrA regulon to a level where it becomes insupportable for efficient cellular cycling in mutant cells that already activate the early S-phase regulon through another, GcrA-independent pathway that can be activated by mutations in *hprK*. Interestingly, further expression of *ctrA* wild-type from vanillate-inducible P*_van_* promoter on the pMT335 plasmid (pMT335-*ctrA*), in the suppressors improves slightly viability of the strains similarly to the Δ*gcrA*::Ω Δ*gcrB* cells (Fig. S5C). However, while expression of *ctrA*Δ*3*Ω, a non-degradable form of the protein (pMT335-*ctrA*Δ*3*Ω) does not impair *WT* viability, the Δ*gcrA*::Ω Δ*gcrB* and suppressors cells tolerate less its expression even in absence of inducer. This result suggests that constitutive expression of CtrA independently of the cell cycle regulation is deleterious to the Δ*gcrA*::Ω Δ*gcrB* cells and further confirm that the suppressors are able to cycle again.

Finally, we introduced an in-frame deletion in *hprK* (Δ*hprK*) and found that this mutation confers a slow-growth phenotype in PYE (Fig. S6A), as described for *S. meliloti*^18^ and the *hprK1.4* point mutant of *C. crescentus*^9^. The Δ*hprK* and *hprK1.4* mutations both lead to hyper-motility on swarm plates (Fig. 4A) and an abundance of G1 cells, particularly in *hprK1.4* cultures, and to a lesser extent in Δ*hprK* cultures (Fig. 4B and S6B). Importantly, transduction of the Δ*gcrA*::Ω deletion into the Δ*hprK* or *hprK1.4* strain yielded colonies after ∼3 days on PYE medium compared to ∼6 days for transduction into *WT* (Fig. 4C). However, the resulting Δ*hprK* Δ*gcrA*::Ω and *hprK1.4* Δ*gcrA*::Ω double mutants showed the same motility/cell cycle defect as the Δ*gcrA*::Ω parent (Fig. 4A, 4B). Thus, even though *hprK1.4* can promote expression of the GcrA-dependent early S-phase regulon, this contribution does not suffice for proper cell cycle (motility) control of Δ*gcrA*::Ω Δ*gcrB* cells to the level seen in the three motility suppressor mutants. Therefore, other mutations are required to allow efficient cycling when GcrA/B are absent.

**Figure 4.**
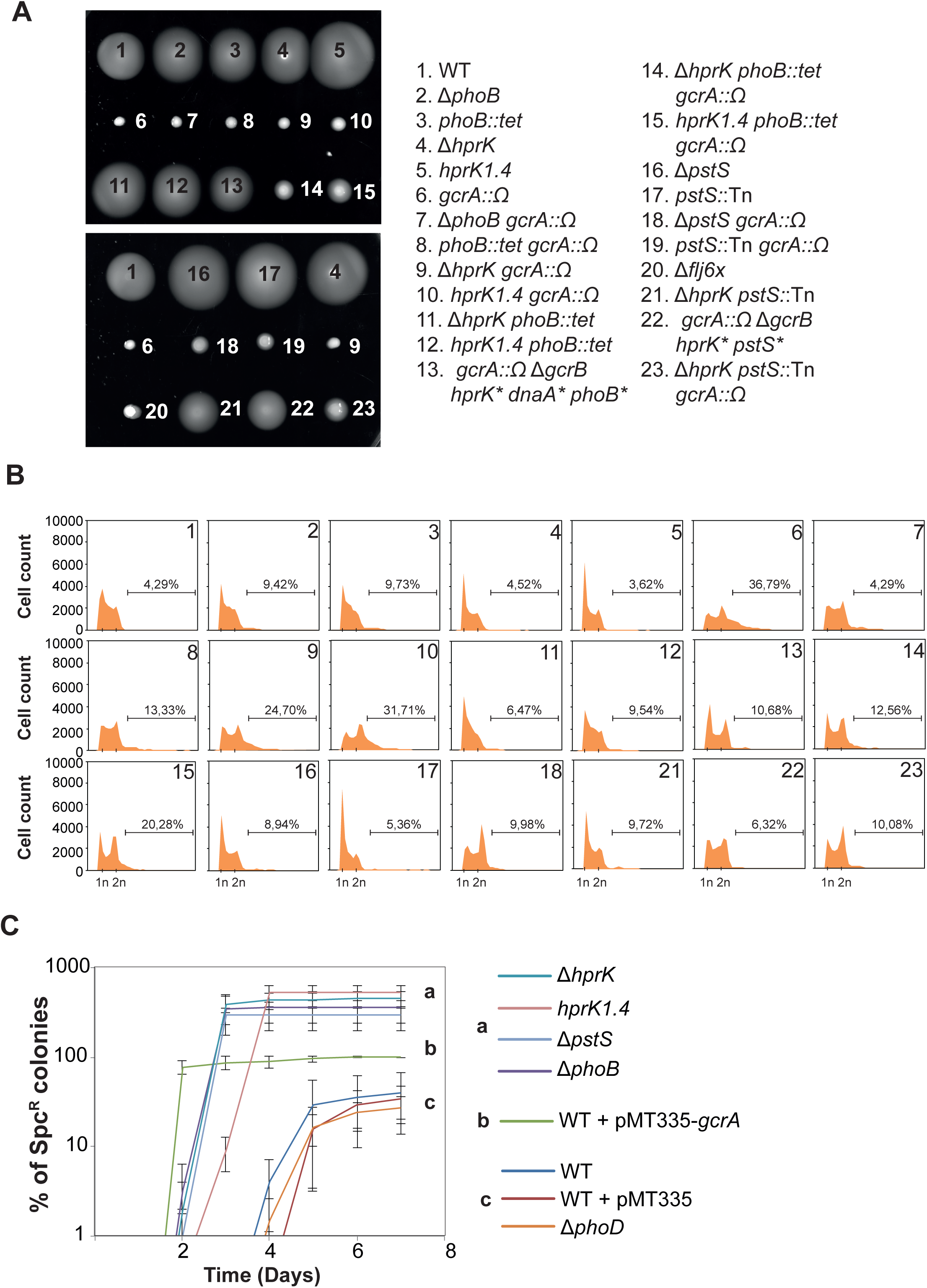
Simultaneous mutations in *hprK* and in the *pho* system are necessary for proper cell cycling (motility) in the absence of GcrA. (**A**) Motility assay on swarm (0.3%) agar and (**B**) DNA content analysis as determined by FACS (FL1-A channel) of *WT*, Δ*fljx6,* Δ*hprK, hprK1.4,* Δ*phoB,* Δ*pstS,* Δ*gcrA*::Ω mutant cells and derivatives during exponential growth in PYE. Percentages of cells containing >2 chromosomes are also indicated. (**C**) Time course of colony appearance following ΦCr30-mediated generalized transduction of Δ*gcrA:*:Ω into different recipient strains. Error bars are standard deviation from three biological replicates. Transductants are scored on PYE media supplemented with spectinomycin (30 µg/mL) and streptomycin (5 µg/mL) to select for the acquisition of the Δ*gcrA::*Ω mutation. Total number of Spc^R^ clones obtained after transduction of cells containing the pMT335-*gcrA* plasmid serves as control and reflects the efficiency of transduction of Δ*gcrA::*Ω.

### Convergence of inorganic phosphate and (p)ppGpp starvation signals on early S-phase promoters

Each of the three motile suppressor mutants harbour a mutation in genes controlling inorganic phosphate uptake in addition to the *hprK^D47E^* mutation. Most bacteria respond to phosphate limitation through a highly conserved signal transduction pathway collectively known as the Pho system^19^. In this pathway, the availability of inorganic phosphate is thought to be sensed by the PhoR-PhoB two-component signalling system that activates the transcription of the high-affinity Pst phosphate uptake transporter in response to phosphate limitation^20^. The first motile suppressor mutant features single missense mutation in the *pstS* gene encoding the periplasmic phosphate binding protein PstS (*pstS^G61S^*) while the second mutant has a missense mutation in the *phoD* gene encoding a secreted alkaline phosphatase PhoD (*phoD^V117A^*), required for the utilization of organic phosphate. The third motile suppressor mutant has two missense mutations in addition to the *hprK^D47E^* mutation: the first one in the *phoB* gene encoding the PhoB response regulator of the PhoR/PhoB two-component system (*phoB^T74A^*), and a second mutation in the *dnaA* gene encoding the bifunctional DnaA replication initiator protein/transcriptional activator (*dnaA^H334R^*).

Several findings indicate that the phosphate starvation response is activated in all three motile suppressor mutants. First, immunoblotting using polyclonal antibodies to the PhoB transcriptional regulator revealed massively elevated steady-state levels of PhoB in all three suppressor mutants compared to *WT* or Δ*gcrA*::Ω Δ*gcrB* parental cells (Fig. 2D). Since PhoB expression regulates its own expression via a promoter upstream of the *pstCAB-phoUB* operon as determined by ChIP-Seq assays using PhoB antibody (Fig. S7A), we infer that the phosphate starvation response is genetically triggered in all three suppressor mutants. Second, the cell division defects are ameliorated (Fig. 5A) and the fraction of G1 cells in the population is augmented (Fig. 5B) when the Δ*gcrA*::Ω Δ*gcrB* parental cells were shifted from medium that has been diluted five-fold with water (PYE/5), a medium known to induce phosphate starvation^21^. Immunoblotting confirmed that the PhoB protein is induced (Fig. 5C) and we found the phosphate starvation promoter probe reporter P*_pstC_*-*lacZ* to be activated in *WT* cells transferred to PYE/5 (Fig. 5D). Moreover, this induction of P*_pstC_*-*lacZ* in PYE/5 was attenuated in Δ*phoB* cells (35% *WT* activity, Fig. 5E) showing that transcriptional activation of P*_pstC_* requires PhoB. By contrast, when we introduced an in-frame deletion in *pstS* (Δ*pstS*) in *WT* cells, we observed the P*_pstC_*-*lacZ* activity to be strongly augmented (336% *WT* activity, Fig. 5F), with a commensurate surge in the levels of PhoB (Fig. 5F). Complementation of the Δ*pstS* mutant with *WT pstS*, but not *pstS^G61S^*, corrected these defects, indicating that the *pstS^G61S^* allele is not functional. In addition, and proving that phosphate starvation response is also required to allow efficient cycling in absence of GcrA/B, we found that wild-type PstS expressed *in trans* (from plasmid pMT335-*pstS*) in Δ*gcrA*::Ω Δ*gcrB hprK^D47E^ pstS^G61S^* motility suppressor cells abolished the motile phenotype (Fig. S8A). In Δ*pstS* cells the chromosomal occupancy of PhoB is globally augmented compared *WT* in ChIP-Seq analyses (Fig. S7A). Bioinformatic analyses of the ChIP-Seq data identified 121 PhoB bound sites (Table S2) based on a five-fold enrichment of PhoB occupancy in either *WT* and/or Δ*pstS* cells (Fig. S7B) *versus* input DNA controls. As expected, the promoters of the *pstS* gene, of the *pstCAB-phoUB* and of *phnCDE* (phosphonates transport system) operons are found in the major targets of PhoB (Fig. S7B, S7C). MEME^22^ analysis predicted, in each of the 98 retained PhoB enriched sites in the Δ*pstS* background, a highly conserved and complex 18 nucleotides (nt) PhoB consensus motif, consisting of two *pho* box repeats (5’-YGTCAYR-3’) separated by a conserved A-rich 4 nt spacer (E-value 2.8e-58) (Fig. S7D). Contrary to what has been described in *C. crescentus*^23^*, pho* boxes seem to adopt a similar architecture as the ones described for *E. coli*, characterized also by two repeats of 7 nt (5’-CTGTCAT-3’) with a 4 nt spacer as well^24^. However, our PhoB ChIP-Seq analysis failed to reveal an overlap between the PhoB and GcrA bound promoters, indicating that the mechanism by which the phosphate starvation response bypasses the Δ*gcrA*::Ω Δ*gcrB* cell cycle defect is PhoB independent or indirect.

**Figure 5.**
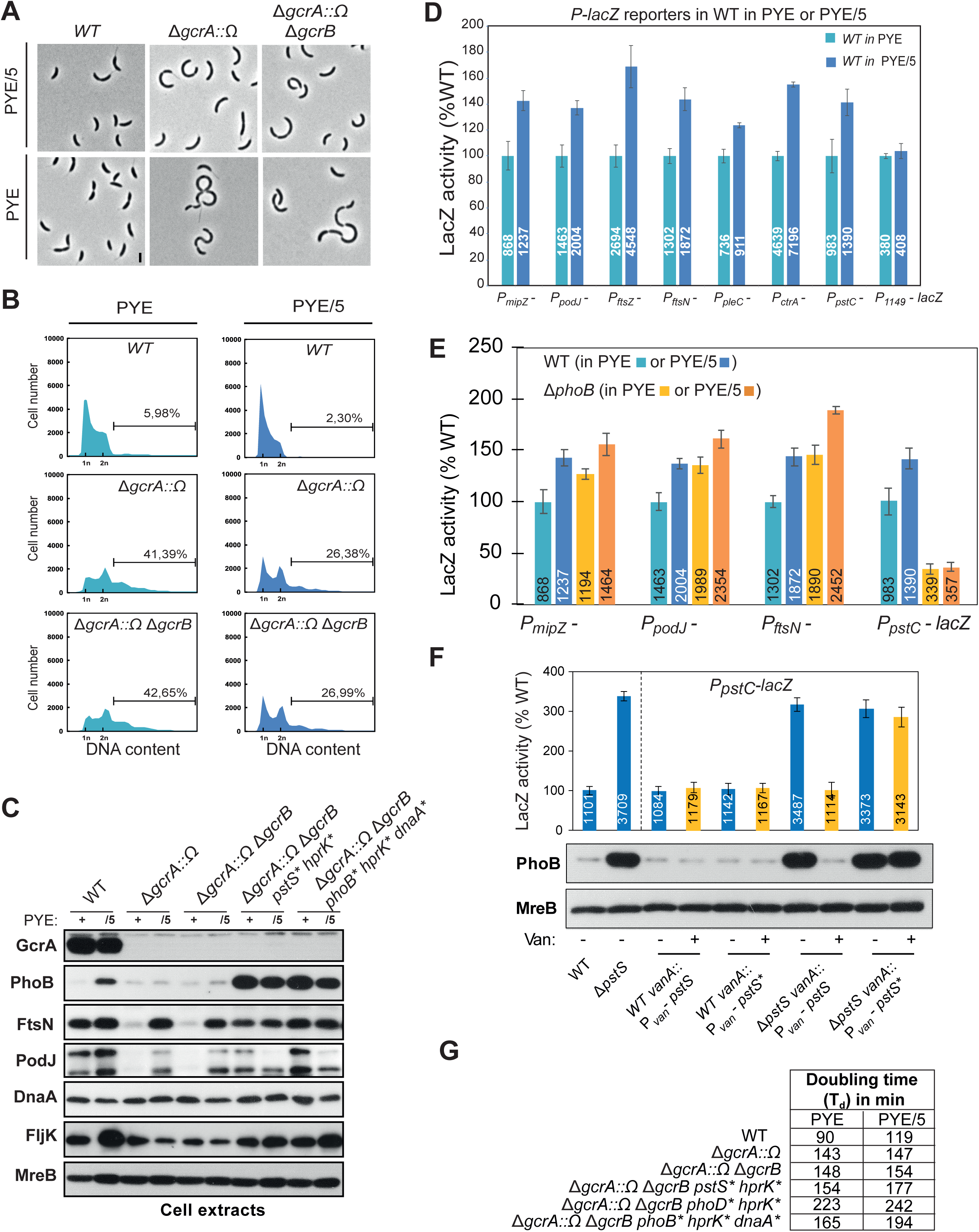
Phosphate starvation rescues S-phase promoter expression in cells lacking GcrA. (**A**) Phase contrast microscopy and (**B**) DNA content analysis as determined by FACS (FL1-A channel) of cells during exponential growth in PYE or PYE/5. Percentages of cells containing >2 chromosomes are also indicated. (**C**) Immunoblots showing steady-state levels of various protein in extracts from *WT*, Δ*gcrA::*Ω, Δ*gcrB* Δ*gcrA::*Ω, Δ*gcrB* Δ*gcrA::*Ω *hprK*\* *pstS*\* and Δ*gcrB* Δ*gcrA::*Ω *hprK*\* *phoB*\**dnaA*\* cells during exponential growth in PYE (+) or 5 hours after shifting to phosphate-limited PYE/5 (/5). Increased steady-state level of PhoB in phosphate-limited conditions serves as positive control. MreB serves as a loading control. (**D**) β-galactosidase analysis measuring the activities of several GcrA-target promoter probe plasmids in *WT* cells during exponential growth in PYE or 5 hours after shifting to phosphate-limited PYE/5. The P*_pstC_*-*lacZ* promoter probe plasmid serves a positive control for the phosphate-limited conditions. The P*_CCNSA_0149_*-*lacZ* (P*_0149_*-lacZ) promoter probe plasmid is GcrA and phosphate starvation independent and thus serves as a negative control. This promoter is not bound by GcrA as determined by previous ChIP-Seq analyses. (**E**) β-galactosidase activities of P*_mipZ_-*, P*_podJ_-*, P*_ftsN_-* or P*_pstC_-lacZ* promoter probe plasmids in *WT* and Δ*phoB* cells grown in PYE to exponential phase or 5 hours after shifting to PYE/5. (**F**) β-galactosidase activities of the P*_pstC_-lacZ* promoter probe plasmid harboring the PhoB controlled *pstC* promoter fused to the promoterless *lacZ* gene in *WT* and Δ*pstS* strains, complemented with *pstS* or *pstS^G61S^* under control of the vanillate-inducible P*_van_*-promoter integrated at the *vanA* locus (pMT384 integrative plasmid), were measured after 5 hours of growth in PYE or PYE supplemented with 50 μM Vanillate. Immunoblotting showing the corresponding PhoB steady-state level in the cells is also indicated. (**D**, **E** and **F**) β-galactosidase activities measured as Miller units are expressed in percentages relative to *WT*. Data are from four independent experiments; error bars are standard deviation.

The fact that PhoB levels are not induced upon shifting Δ*gcrA*::Ω or Δ*gcrA*::Ω Δ*gcrB* cells from PYE to PYE/5 (Fig. 5C) provides further evidence that signaling via the PhoBR two-component system is not required to induce the GcrA-regulon independently of GcrA. To further illuminate the link between inorganic phosphate control and early S-phase (GcrA-dependent) promoter firing, we measured the activity of several *lacZ*-based promoter probe reporters to GcrA-bound promoters, before and after shifting *WT* cells from PYE to PYE/5 (Fig. 5D). We observed an increase in β-galactosidase activity for each tested GcrA-target promoter, indicating that the GcrA regulon is induced when cells are starved in PYE/5. However, when we measure the activity of the P*ftsN* promoter in Δ*spoT* cells shifted in PYE/5 (Fig. 3D), we observed a decrease in β-galactosidase activity suggesting that transcription of S-phase promoters in PYE/5 is the result of both phosphate starvation and (p)ppGpp induction. Consistent with the suppressor analysis described above, we discovered that the activity of these GcrA-target promoters is also augmented in Δ*phoB* cells and Δ*pstS* cells grown in PYE (Fig. 5E, S4B). Since Δ*phoB* cells induce the early S-phase regulon even further under nutrient depletion (in PYE/5, Fig. 5E), it is evident that cells experience an additional stress other than phosphate deprivation in PYE/5, again consistent with the multifactorial genetic alterations in the Δ*gcrA*::Ω Δ*gcrB* motility suppressor mutants.

We draw two important conclusions from these experiments. First, cells reprogram early S-phase transcription in diluted medium (PYE/5) as a result of a combined phosphate starvation and the (p)ppGpp induction response (Fig. 3D). Second, early S-phase transcriptional reprogramming can also be achieved in nutrient replete conditions by (at least two) genetic modifications, including a mutation in the phosphate starvation pathway and a mutation in the (p)ppGpp activation pathway, that can restore cellular cycling to Δ*gcrA*::Ω Δ*gcrB* cells. To further support the necessity of this two combined genetic modifications, we confirmed that the motility phenotype could only be restored after transduction of the Δ*gcrA*::Ω mutation into mutants in which both pathways are simultaneously mutated: i.e. cells having mutations in *hprK* (either Δ*hprK* or *hprK1.4*) and the phosphate starvation pathway (either Δ*phoB::tet* or *pstS::Tn*) (Fig. 4A).

### Conserved induction of cell cycle promoters by phosphate starvation

Matching findings for a multifactorial suppression mechanism also emerged from motility suppressor analyses of Δ*gcrA*::Ω *ccrM*::Tn cells. We isolated 5 motile Δ*gcrA*::Ω *ccrM*::Tn suppressor mutants on swarm agar (Fig. 6A) and sequenced their genomes (Fig. 6B). Four out of the five mutants have two missense mutations: one in *hprK^D47E^*, similar to the one isolated in the Δ*gcrA*::Ω Δ*gcrB* double mutant along with another one in the *pst* (phosphate transporter) system in *pstB* (*pstB*^R234W^, *pstB*^S236P^ and *pstB*^stop275R^) and *pstC* (*pstC*^L379P^). By contrast in the last suppressor mutant a *pstC* (*pstC*^S446P^) mutation is also present, but the *hprK* gene is not altered. Instead, we found a missense in the gene encoding the E1 component of the PTS system that regulates SpoT (*ptsP*^A130D^). Thus, again the genetic constitution of the suppressor mutants 1-4 and suppressor mutant 5 shows that HprK-PtsP-SpoT along with the phosphate starvation pathway need to be jointly modified in order to permit cycling of Δ*gcrA*::Ω *ccrM*::Tn cells (Fig. S2B).

**Figure 6.**
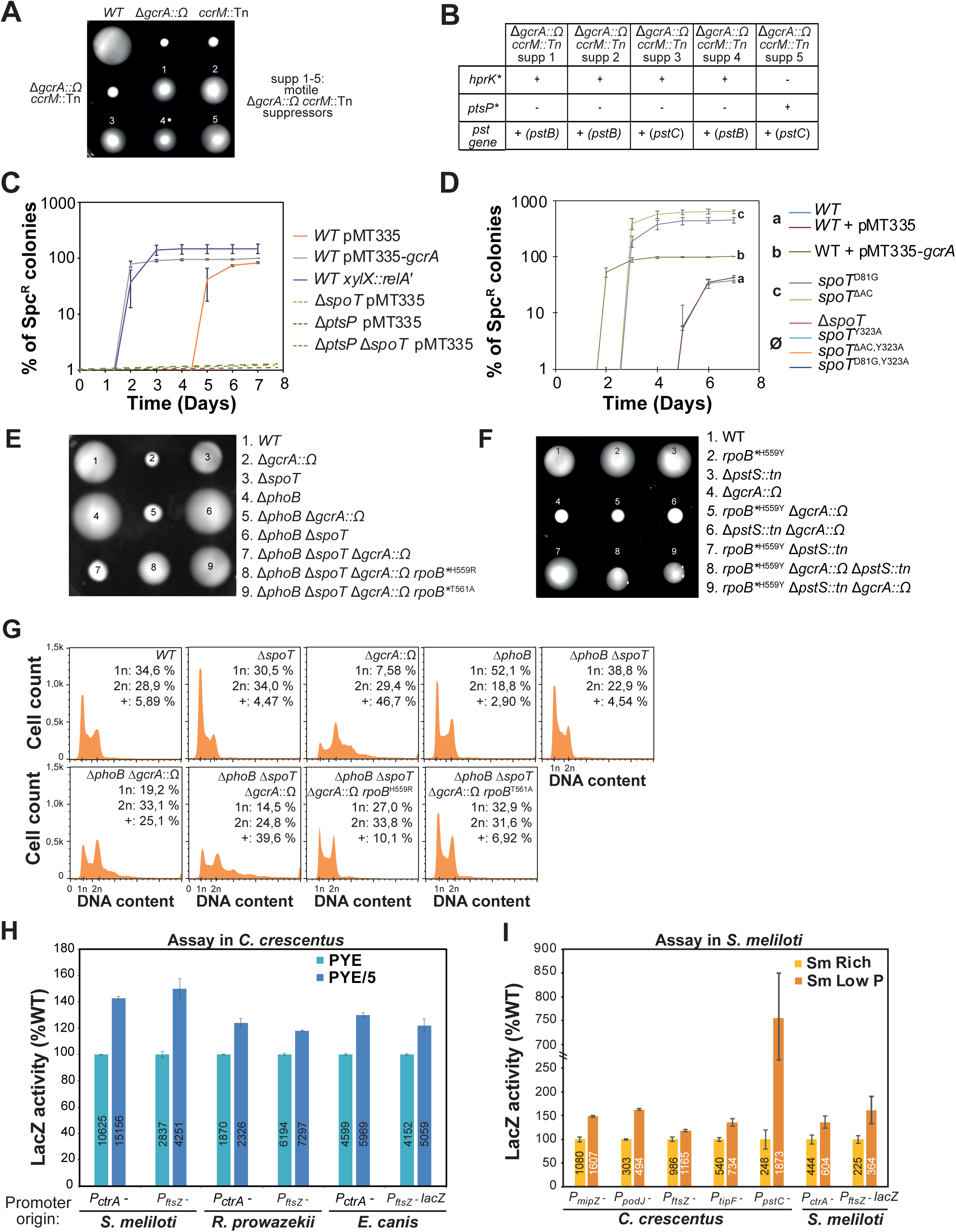
The PTS system and (p)ppGpp alarmone compensate for the loss of GcrA independently of GANTC promoter methylation. (**A**) Motility assays on swarm (0.3%) agar of *WT*, Δ*gcrA::*Ω and *ccrM::*Tn single mutant cells, Δ*gcrA::*Ω *ccrM::*Tn double mutant cells and five isolated spontaneous motility suppressor mutants derived from Δ*gcrA::*Ω *ccrM::*Tn double cells. (**B**) Whole genome sequencing of the motility suppressors revealed a combination of two point mutations: (i) one in the PTS system pathway (either *hprK^D47E^* (1 to 4) or *ptsP^A130D^* (5)) and (ii) a second in the *pho* system (either *pstB^R234W^* (1), *pstB^S236P^* (3), *pstB^stop275R^* (4), *pstC^L379P^* (2) or *pstC^S446P^* (5)). (**C** and **D**) Time course of colony appearance following ΦCr30-mediated generalized transduction of Δ*gcrA::*Ω into various recipient cells, selected on PYE plates supplemented with spectinomycin (30 µg/mL) and streptomycin (5 µg/mL) to select for the Δ*gcrA::*Ω mutation. The total number of Spc^R^ clones obtained after transduction of *WT* cells containing the pMT335-*gcrA* plasmid serves as control and reflects the efficiency of transduction of Δ*gcrA::*Ω. (**E** and **F**) Motility assays on swarm (0.3%) agar of indicated strains. (**G**) DNA content analysis as determined by FACS (FL1-A channel) of cells grown exponentially in PYE. Percentages of cells containing 1n, 2n and >2 chromosomes are also indicated. (**H** and **I**) β-galactosidase activities of P*_mipZ_-*, P*_podJ_-*, P*_ftsZ_-,* P*_tipF_-* or P*_pstC_-lacZ* fusions from *C. crescentus,* P*_ctrA_-* and P*_ftsZ_-lacZ* fusion of promoters from different alpha-proteobacteria (*Sinorhizobium meliloti*, *Rickettsia Prowazekii* and *Ehrlichia canis*) were determined in *WT C. crescentus* (**H**) during exponential growth in PYE or 5 hours after shifting to PYE/5, or in *WT S. meliloti* cells (**I**) grown in Sm Rich or Sm Low P (low phosphate) medium as indicated. Sm corresponds to a specific *S. meliloti* growth media. P*_pstC_*-promoter serves a positive control for the phosphate-limited conditions. β-galactosidase activities measured as Miller units are expressed in percentages relative to *WT*. Data are from four independent experiments; error bars are standard deviation.

The isolation of the mutation in *ptsP* strongly supports the notion that (p)ppGpp can promote efficient cycling in cells lacking GcrA, provided the phosphate starvation response is activated. We recreated this genetic context by transducing the Δ*gcrA*::Ω mutation into Δ*pstS* cells and Δ*phoB* cells harbouring the xylose-inducible heterologous (p)ppGpp-synthase RelA’ from *E. coli* integrated at the chromosomal *xylX* locus of *C. crescentus*.

Motility assays conducted on swarm agar indicate a substantial increase in motility upon induction of RelA’ with xylose, while substantially less motility was observed in the presence of glucose which represses RelA’ expression (Fig. S8B). As motility is an indicator for cellular cycling of *C. crescentus* cells, we surmise that cellular cycling was in large ameliorated in these genetically modified Δ*gcrA*::Ω cells, a conclusion that was upheld by FACS analyses (Fig. S9). Conversely, we expected that a failure to produce (p)ppGpp should have an adverse effect on cycling of Δ*gcrA*::Ω cells. In fact we observed that it was impossible to transduce the Δ*gcrA*::Ω mutation into Δ*spoT* and Δ*ptsP* single mutant cells or into Δ*spoT* Δ*ptsP* double mutant cells (Fig. 6C and 6D). In addition, it was also impossible to transduce the Δ*gcrA*::Ω mutation into *spoT^syn^*\* (*spoT^Y323A^*) variants whose (p)ppGpp synthase activity is abolished (Fig. 6D). Importantly, transduction of the Δ*gcrA*::Ω deletion into *spoT^hyd^*\* variants (*spoT^ΔAC^* and *spoT^D81G^*) defective in (p)ppGpp hydrolysis yielded colonies after ∼3 days on PYE medium compared to ∼6 days for transduction into *WT* (Fig. 6D), thus confirming that (p)ppGpp signaling is required for efficient growth in the absence of GcrA.

The need for (p)ppGpp for growth can be bypassed when phosphate starvation response is induced, i.e. in Δ*phoB* Δ*spoT* cells it was possible to inactivate *gcrA* by transduction of the Δ*gcrA*::Ω mutation. However, motility assays on swarm agar shows that the resulting triple mutant cells are poorly motile akin to the Δ*gcrA*::Ω cells (Fig. 6E) and thus do not cycle properly (Fig. S8C). Importantly, genome sequencing of two motility suppressors isolated from Δ*phoB* Δ*spoT* Δ*gcrA*::Ω cells (Fig. 6E) revealed that each mutant had a missense mutation in the *rpoB* gene: encoding either RpoB^H559R^ or RpoB^T561A^. Thus, phosphate-starvation and (p)ppGpp signaling converge to the same cellular GcrA-dependent transcriptional pathway on RNA polymerase, as (p)ppGpp-blind mutation in the same residue of RpoB (H559Y) augment transcription from GcrA-dependent promoters (Fig. S8E) and allows to restore motility and chromosome content in the triple *rpoB*^H559Y^* Δ*pstS*::tn Δ*gcrA*::Ω mutant cells (Fig. 6F and 6G). Finally, we asked if this mechanism of induction of early S-phase promoters is conserved in alpha-proteobacteria. To this end, we cloned the orthologous promoters that are bound by GcrA in *C. crescentus* into *lacZ*-based promoter probe vectors and tested them for induction in *C. crescentus WT* cells in PYE and PYE/5 (Fig. 6H). We observed that the promoters upstream of the *ftsZ* and *ctrA* coding sequences in *Sinorhizobium meliloti*, *Rickettsia prowazekii* and *Ehrlichia canis* were all induced in PYE/5 relative to PYE. Conversely, we probed for activity of several GcrA-target promoters from *C. crescentus* in *S. meliloti* cells grown in rich medium or under phosphate limitation^25^ (Fig. 6I). These heterologous reporters of GcrA were also induced in phosphate starved *S. meliloti* cells. Lastly, we observed that the native *S. meliloti ftsZ* and *ctrA* promoter reporters were similarly induced in phosphate starved *S. meliloti* cells, indicating that a comparable interplay between phosphate starvation and cell cycle control also exists in α-proteobacteria other than *C. crescentus*.

### GcrA recruits RNAP to promoters and regulation by phosphate starvation

To illuminate the mechanism by which the GcrA regulon is transcriptionally activated by phosphate starvation, we conducted quantitative (q)ChIP experiments in *C. crescentus* cells grown in PYE versus PYE/5 using polyclonal antibodies to GcrA and a monoclonal antibody to the housekeeping sigma factor RpoD/σ^70^ to quantify their abundance at the *ftsN* and *pstC* promoters (P*_ftsN_* and P*_pstC_*, Fig. 7) relative to a neutral site such as the replication terminus (*ter*). As expected from previous ChIP-Seq experiments in which we had used antibodies to GcrA to precipitate chromatin from *WT* cells, GcrA associates very efficiently with P*_ftsN_*, whereas it is nearly undetectable at P*_pstC_*. Similarly, only background precipitations of chromatin from Δ*gcrA*::Ω cells was observed using antibodies to GcrA. Importantly, our qChIP experiment showed that GcrA indeed recruits σ^70^ to P*_ftsN_*, contrary to the model proposed by Haakonsen *et al*. in which GcrA is positioned at the promoter by σ^70^. In fact, we observed a drastic (approximately 20-fold) reduction in σ^70^ occupancy at P*_ftsN_* in Δ*gcrA*::Ω cells compared to *WT* cells and reduced P*_rpoD/σ70_*-*lacZ* activity (68% compared to *WT*) in cells depleted for GcrA (Fig. S10). We also observed that σ^70^ occupancy at P*_pstC_* increases substantially in Δ*pstS* mutant cells grown in PYE compared to control cells, consistent with the induction of the P*_pstC_*-*lacZ* reporter. Importantly, this recruitment of σ^70^ to P*_pstC_* is attenuated in Δ*phoB* cells, indicating that both GcrA and PhoB play crucial roles in transcription initiation through recruitment of σ^70^ to the promoter when they are active.

**Figure 7.**
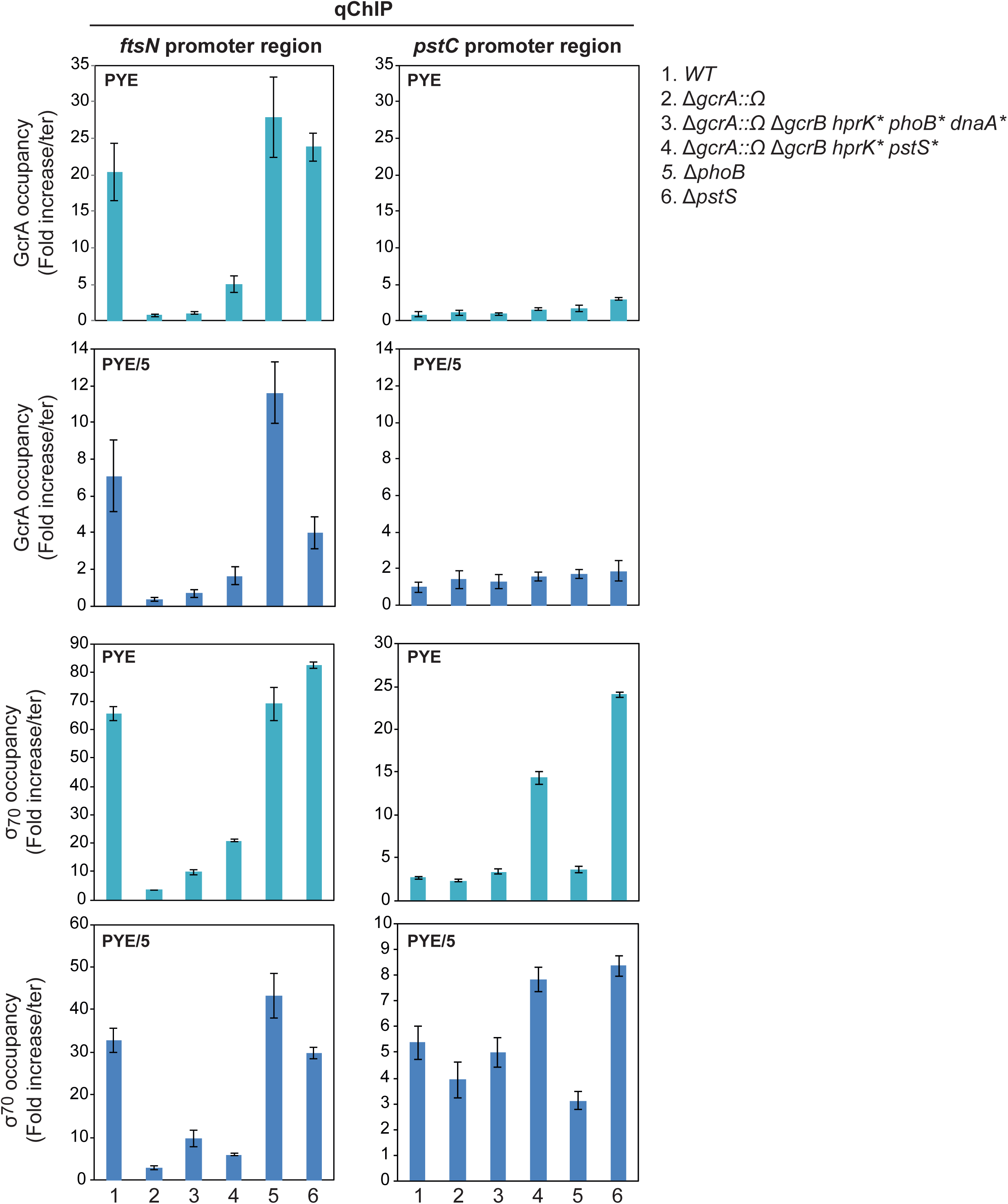
Promoter occupancy of σ^70^ at P*_ftsN_* is controlled by GcrA and phosphate starvation rescues *ftsN* expression in a GcrA/σ^70^ independent manner. Chromatin immunoprecipitation followed by quantitative real time PCR (qChIP) performed on *WT*, Δ*gcrA::*Ω, Δ*gcrB* Δ*gcrA::*Ω *hprK*\* *phoB*\**dnaA*\*, Δ*gcrB* Δ*gcrA::*Ω *hprK*\* *pstS*\*, Δ*phoB* and Δ*pstS* cells. The qChIP data shows the GcrA and the RpoD/σ^70^ occupancy at the promoter of *ftsN* and *pstC* (relative to a neutral site such as the replication terminus (*ter*)) during exponential growth in PYE (light blue) or 5 hours after shifting to PYE/5 (dark blue). The *pstC* promoter region serves as a positive control for phosphate starvation condition and for σ^70^ occupancy at this promoter as PhoB binds to region 4 of σ^70^ in *E. coli.* Data are from 3 independent experiments; error bars are standard deviation.

While qChIP experiments also did reveal a small increase in σ^70^ recruitment to GcrA target promoters when Δ*gcrA*::Ω cells are grown in PYE/5, the increase does not seem commensurate with the increase in promoter firing under phosphate starvation ((p)ppGpp-inducing conditions). As (p)ppGpp has also been predicted to control RNA polymerase (RNAP) in α-proteobacteria, we hypothesize that (p)ppGpp promotes promoter escape or the transcription elongation phase in Δ*gcrA*::Ω suppressor mutants.

## Discussion

The transcriptional cell cycle regulator GcrA that works together with the CcrM adenine methyltransferase to preferentially activate transcription at methylated promoters fire in S-phase, to give a burst of expression of highly conserved cell division (FtsN, MipZ, FtsZ), polarity (PodJ, TipF, ZitP) and regulatory proteins (CtrA) that are required for cell cycle progression and polar development in alpha-proteobacteria. Several alpha-proteobacteria including *C. crescentus* and *S. meliloti* encode GcrA and CcrM proteins, however, the obligate intracellular rickettsial lineages do not. Our mutants sustain cellular cycling and polar development in the absence of GcrA, presumably because firing of GcrA/CcrM-dependent S-phase promoters is bypassed by the suppressor mutations activating phosphate starvation and (p)ppGpp signalling. The fact that both GcrA-dependent promoters from either *C. crescentus* or *S. meliloti* are active in the heterologous host upon phosphate starvation (Fig. 6H and 6I), indicates that the induction of S-promoter activity in the absence of GcrA is conserved. The rickettsial cell cycle is supported without a GcrA/CcrM system, and the finding that *ftsZ* and *ctrA* promoter sequences from two different members of the *Rickettsiales* to respond in *WT C. crescentus* with increase in activity upon phosphate starvation, offers an explanation to allow the *Rickettsiales* to lose genes encoding the GcrA/CcrM module during evolution, while sustaining a (slow) cell cycle.

Phosphate starvation and (p)ppGpp signalling can support transcription of S-phase promoters and cellular cycling even when GcrA and CcrM are both absent as indicated that mutations in these pathways also surface in motile *ccrM::*Tn Δ*gcrA*::Ω suppressor mutants. Consistent with the conclusion that the bypass pathway acts independently of methylation, we show that GANTC mutated *ftsN* promoter also responds to increased levels of (p)ppGpp (Fig. 3C). We previously demonstrated that upon entry in stationary phase the GcrA-regulon is induced in a SpoT dependent manner^17, 26^ (Fig. 3B) suggesting a role of (p)ppGpp in transcription activation of the early S-phase promoters. Moreover, recent motility suppressor screen analysis of the Δ*phoB* Δ*spoT* mutant leads to the identification of mutation in the *rpoB* gene, encoding the β-subunit of the RNA polymerase. Interestingly, such *rpoB\** mutations increase the motility of the *WT C. crescentus* strain and increases the promoter activity of early S-phase genes similarly to ectopic (p)ppGpp induction but does not suppress the motility defect of the Δ*gcrA*::Ω mutant cells. This suppressor mutation further confirms our precedent findings that first induction of (p)ppGpp leads to an induction of GrcA-dependent regulon and signals via the RNAP and second that both phosphate starvation induced mechanism and (p)ppGpp dependent effect on the RNAP are necessary to help the Δ*gcrA*::Ω mutant cells to restore cell cycle (Fig. 6F). Importantly, mutation in the *rpoB* or *rpoC* genes, encoding the β and β’ subunit of the RNAP respectively, can suppress the defect of the *S. meliloti rel* mutant deficient for the production of (p)ppGpp^27^. Moreover, similarly to what has been observed in *C. crescentus*, (p)ppGpp accumulation in *S. meliloti*^15, 28^ also induces a G1-arrest suggesting a conserved role in (p)ppGpp cell cycle regulation.

(p)ppGpp alarmone clearly plays a role in S-phase promoter activation even under normal conditions as GcrA becomes indispensable in a Δ*spoT* background similarly to Δ*ptsP* or Δ*ptsP* Δ*spoT* mutant. This result implicates the PTS system in the control of (p)ppGpp synthesis. However, it was possible to transduce the *gcrA::Ω* deletion in the Δ*phoB* Δ*spoT* mutants cells, explaining why both mutation (PTS and *pho*) are necessary to allow restoration of the cell cycle and suggesting that phosphate starvation does not act at the level of transcription through induction of (p)ppGpp, at least through the only known (p)ppGpp synthase in *C. crescentus*.

The mechanism by which phosphate starvation and environmental cues cell cycle dependant transcription is unknown, but the recruitment of σ^70^ to the *ftsN* promoter increases in the Δ*gcrA*::Ω suppressor mutants versus the Δ*gcrA*::Ω parent, implicating the phosphate starvation response in the recruitment of σ^70^ to S-phase promoters. The strong reduction of σ^70^ occupancy in Δ*gcrA*::Ω cells also shows that σ^70^ promoter recruitment depends on GcrA. It has been proposed that GcrA binds to all active σ^70^ promoters via its interaction with σ^70^ domain 2 and the RNAP^4, 5^, but the dependency on RNAP/σ^70^ was never demonstrated *in vivo.* Importantly, our ChIP analyses and this indicate that GcrA does recruit or at least stabilize the σ^70^•RNAP holoenzyme. In support of this, we previously showed that ChIP-seq experiments with antibodies to GcrA detect preferential binding of GcrA at a subset of promoters containing methylated GANTC sites, not at all σ^70^ promoters. On the basis of our previous findings and those described by Haakonsen *et al*, along with the data reported here, we propose a revised model in which GcrA binds σ^70^ and is recruited to all σ^70^-dependent promoter, including S-phase promoters that harbour a methylation mark. When GcrA recognizes the methylation mark, its interaction with target DNA is stabilized which also stabilizes σ^70^•RNAP at that site and allows for promoter isomerization and firing. In the absence of the GcrA and CcrM-dependent activation convergent nutrient starvation signalling allows for increased σ^70^•RNAP abundance at the level of initiation, but also improved elongation mediated by (p)ppGpp.

In sum the genetic and likely evolutionary plasticity of S-phase promoter activation or at least transcription (initiation and/or elongation) by σ^70^•RNAP in the alpha-proteobacteria allows for sufficient temporal precision to sustain cellular cycling in the absence of GcrA when phosphate starvation and (p)ppGpp signalling are jointly activated.

## Material and Methods

### Growth conditions

*Caulobacter crescentus* NA1000^29^ and derivatives were cultivated at 30°C in peptone yeast extract (PYE) rich medium. 1.5% agar was added into PYE plates and motility was assayed on PYE plates containing 0.3% agar. Antibiotic concentrations used for *C. crescentus* include kanamycin (solid: 20 µg/mL; liquid: 5 µg/mL), tetracycline (1 µg/mL), spectinomycin (liquid: 25 µg/mL), spectinomycin/streptomycin (solid: 30 and 5 µg/mL, respectively), gentamycin (1 µg/mL) and nalidixic acid (20 µg/mL). When needed, final concentration of D-xylose (0.3%), sucrose (0.3%), glucose (0.2%), vanillate (50μM) and IPTG (0.1mM) were added. Swarmer cell isolation, electroporation, biparental mating and bacteriophage φCr30-mediated generalized transduction were performed as described in^30–32^.

*Sinorhizobium meliloti* Rm2011^33^ and derivatives were cultivated at 30°C in Luria broth (LB) rich medium supplemented with CaCl_2_ 2.5 mM and MgSO_4_ 2.5 mM. For S. *meliloti*, nalidixic acid and tetracycline antibiotics were used at 8 and 10 μg/mL, respectively.

*Escherichia coli* S17-1^30^ and EC100D (Epicentre Technologies, Madison, WI, USA) were cultivated at 37°C in LB).

### Phosphate starvation conditions

Rich medium overnight cultures (PYE) of *C. crescentus* were harvested and washed 3 times with PYE/5 (5-fold diluted PYE except for MgSO_4_ 1 mM and 1mM CaCl_2_, supplemented with 0.2% glucose)^21^, and then restarted in PYE/5 medium for 5h at 30°C.

Rich medium overnight cultures (LB+ CaCl_2_+MgSO_4_) of *S. meliloti* were harvested and washed 3 times with MOPS-minimal medium (40 mM morpholinopropanesulfonic acid (MOPS), 20 mM KOH, 20 mM NH_4_Cl, 2 mM MgSO_4_, 100 mM NaCl, 1.2 mM CaCl_2_, 0.3 mg/ml biotin and 15mM glucose)^25^, and then restarted in MOPS-minimal medium for 5h at 30°C.

### Bacterial strains, plasmids and oligonucleotides

Bacterial strains, plasmids and oligonucleotides used in this study are listed and described in tables below.

### Plasmid constructions

#### pNPTS138*-ΔphoB*

The plasmid construct used for *phoB* (*CCNA_00296*) deletion was made by PCR amplification of two fragments. The first, a 444 bp fragment flanked by an *Eco*R1 site at the 5’end and a *Bam*HI at the 3’end (nt 309915-310359, amplified using primers delphoB_1-EcoRI and delphoB_1-BamHI), encompasses the predicted start codon. The second 471 bp fragment, flanked by a *Bam*HI site at the 5’end and a *Hind*III site at the 3’end (nt 311064- 311535, amplified using primers delphoB_2-BamHI and delphoB_2-HindIII), harbors the last 3 bp of the *phoB* coding sequence and extends 469 bp downstream of the gene. These two fragments were first digested with appropriate restriction enzymes and then triple ligated into pNTPS138 (M.R.K. Alley, unpublished) that had been previously restricted with *Eco*RI and *Hin*dIII.

#### pNPTS138-Δ*pstS*

The plasmid construct used for *pstS* (*CCNA_01583*) deletion was made by PCR amplification of two fragments. The first, a 485 bp fragment flanked by an *Eco*R1 site at the 5’end and a *Bam*HI at the 3’end (nt 1699541 -1700025, amplified using primers delpstS_1- EcoRI and delpstS_1-BamHI), encompasses the upstream region of *pstS* and extends to 21 pb downstream of the predicted start codon. The second 506 bp fragment, flanked by a *Bam*HI site at the 5’end and a *Hind*III site at the 3’end (nt 1698030-1698535, amplified using primers delpstS_2-BamHI and delpstS_2-HindIII), harbors the last 15 bp of the *pstS* coding sequence and extends 491 bp downstream of the gene. These two fragments were first digested with appropriate restriction enzymes and then triple ligated into pNTPS138 (M.R.K. Alley, unpublished) that had been previously restricted with *Eco*RI and *Hin*dIII.

#### pNPTS138-Δ*phoD*

The plasmid construct used for *phoD* (*CCNA_01636*) deletion was made by PCR amplification of two fragments. The first, a 728 bp fragment flanked by an *Eco*R1 site at the 5’end and a *Bam*HI at the 3’end (nt 1753493 -1754220, amplified using primers delphoD_1-EcoRI and delphoD_1-BamHI), encompasses the *phoD* predicted start codon. The second 564 bp fragment, flanked by a *Bam*HI site at the 5’end and a *Hind*III site at the 3’end (nt 1755794-1756357, amplified using primers delphoD_2-BamHI and delphoD_2-HindIII), harbors the last 11 bp of the *phoD* coding sequence and extends 553 bp downstream of the gene. These two fragments were first digested with appropriate restriction enzymes and then triple ligated into pNTPS138 (M.R.K. Alley, unpublished) that had been previously restricted with *Eco*RI and *Hin*dIII.

#### pNPTS138-*ΔhprK*

The plasmid construct used for *hprK* (*CCNA_00239*) deletion was made by PCR amplification of two fragments. The first, a 587 bp fragment flanked by an *Eco*R1 site at the 5’end and a *Xba*I at the 3’end (nt 254225 -254811, amplified using primers delhprK_1-EcoRI and delhprK_1-XbaI), encompasses the upstream region of *hprK* and extends to 4 pb downstream of the predicted start codon. The second 573 bp fragment, flanked by a *Xba*I site at the 5’end and a *Hind*III site at the 3’end (nt 255238-255810, amplified using primers delhprK_2-XbaI and delhprK_2-HindIII), harbors the last 5 bp of the *hprK* coding sequence and extends 568 bp downstream of the gene. These two fragments were first digested with appropriate restriction enzymes and then triple ligated into pNTPS138 (M.R.K. Alley, unpublished) that had been previously restricted with *Eco*RI and *Hin*dIII.

#### pNPTS138-*spoT-*us

PCR was used to amplify a DNA fragment upstream of the *CCNA_01622* ORF. A 902 bp fragment (nt 1741409-1742310, NA1000 genome coordinates, flanked by an *Eco*RI site at the 5’end and a *Hind*III at the 3’end) was amplified using primers spoT-us-1-EcoRI and spoT-us-2-HindIII. The PCR fragment was digested with *Eco*RI/*Hind*III, then ligated into pNPTS138 restricted with *Eco*RI and *Hind*III.

#### pMT384-*pstS*

The *pstS* (*CCNA_01583*) coding sequence was PCR amplified from NA1000 with primers NdeI-*pstS* and EcoRI-*pstS* (with *Nde*I site overlapping the start codon and with *Eco*RI site flanking the stop codon) and cloned into pMT384 restricted with *Nde*I and *Eco*RI.

#### pMT384-pstS^G61S^

The *pstS^G61S^* (*CCNA_01583*) coding sequence was PCR amplified from NA1000 Δ*gcrB* Δ*gcrA*::Ω *hprK*^D47E^ *pstS*^G61S^ with primers NdeI-*pstS* and EcoRI-*pstS* (with *Nde*I site overlapping the start codon and with *Eco*RI site flanking the stop codon) and cloned into pMT384 restricted with *Nde*I and *Eco*RI.

#### pMT335-*pstS*

The *pstS* (*CCNA_01583*) coding sequence was PCR amplified from NA1000 with primers NdeI-*pstS* and EcoRI-*pstS* (with *Nde*I site overlapping the start codon and with *Eco*RI site flanking the stop codon) and cloned into pMT335 restricted with *Nde*I and *Eco*RI.

#### pMT335-pstS^G61S^

The *pstS^G61S^* (*CCNA_01583*) coding sequence was PCR amplified from NA1000 Δ*gcrB* Δ*gcrA*::Ω *hprK*^D47E^ *pstS*^G61S^ with primers NdeI-*pstS* and EcoRI-*pstS* (with *Nde*I site overlapping the start codon and with *Eco*RI site flanking the stop codon) and cloned into pMT335 restricted with *Nde*I and *Eco*RI.

#### pMT335-*hprK*

The *hprK* (*CCNA_00239*) coding sequence was PCR amplified from NA1000 with primers NdeI-*hprK* (with *Nde*I site overlapping the start codon) and XbaI-*hprK* (with *XbaI* site flanking the stop codon) and cloned into pMT335 restricted with *Nde*I and *Xba*I.

#### pMT335-hprK^D47E^

The *hprK^D47E^* (*CCNA_00239*) coding sequence was PCR amplified from NA1000 Δ*gcrB* Δ*gcrA*::Ω *hprK*^D47E^ *dnaA*^H334R^ *phoB*^T74A^ with primers NdeI-*hprK* (with *Nde*I site overlapping the start codon) and XbaI-*hprK* (with *XbaI* site flanking the stop codon) and cloned into pMT335 restricted with *Nde*I and *Xba*I.

#### pMT335-*hprK1.4*

The *hprK1.4* (*CCNA_00239*) coding sequence was PCR amplified from NA1000 *hprK1.4*^9^ with primers NdeI-*hprK* (with *Nde*I site overlapping the start codon) and XbaI-*hprK* (with *XbaI* site flanking the stop codon) and cloned into pMT335 restricted with *Nde*I and *Xba*I.

#### pMT335-*ctrA*

The *ctrA* (*CCNA_03130*) coding sequence was PCR amplified from NA1000 with primers NdeI-*ctrA* (with *Nde*I site overlapping the start codon) and EcoRI-*ctrA* (with *Eco*RI site flanking the stop codon) and cloned into pMT335 restricted with *Nde*I and *Eco*RI.

#### pMT335-*ctrA*Δ*3*Ω

The *ctrA*Δ*3*Ω (*CCNA_03130*) coding sequence was PCR amplified from NA1000 with primers NdeI-*ctrA* (with *Nde*I site overlapping the start codon) and EcoRI-*ctrAΔ3Ω* (with *Eco*RI site flanking the stop codon) and cloned into pMT335 restricted with *Nde*I and *Eco*RI.

#### placZ290-PpstC

The *pstC* promoter region (-454 to +123 relative to the ATG) was PCR-amplified using ppstC-EcoRI and ppstC-XbaI primers using NA1000 chromosomal DNA as a template. This fragment was digested by appropriate enzymes and cloned into a *Eco*RI*-Xba*I-digested p*lacZ*290 promoter probe vector.

#### placZ290-PrpoD

The *rpoD* promoter region (-695 to +28 relative to the ATG) was PCR-amplified using prpoD-EcoRI and prpoD-XbaI primers using NA1000 chromosomal DNA as a template. This fragment was digested by appropriate enzymes and cloned into a *Eco*RI*-Xba*I-digested p*lacZ*290 promoter probe vector.

#### PlacZ290-PEcctrA

A synthetic fragment (DNA2.0 Inc, Menlo Park, CA, USA) encoding the promoter region of the *Ehrlichia canis ctrA* gene was digested by appropriate enzymes and cloned into p*lacZ290* (*Eco*RI/*Xba*I).

#### PlacZ290-PEcftsZ

A synthetic fragment encoding the promoter region of the *Ehrlichia canis ftsZ* was digested by appropriate enzymes and cloned into p*lacZ290* (*Eco*RI/*Xba*I).

#### PlacZ290-PRpctrA

A synthetic fragment encoding the promoter region of the *Rickettsia Prowazekii ctrA* was digested by appropriate enzymes and cloned into p*lacZ290* (*Eco*RI/*Xba*I).

#### PlacZ290-PRpftsZ

A synthetic fragment encoding the promoter region of the *Rickettsia Prowazekii ftsZ* was digested by appropriate enzymes and cloned into p*lacZ290* (*Eco*RI/*Xba*I).

#### PlacZ290-PRm2011ctrA

A synthetic fragment encoding the promoter region of the *Sinorhizobium meliloti ctrA* was digested by appropriate enzymes and cloned into p*lacZ290* (*Eco*RI/*Xba*I).

#### PlacZ290-PRm2011ftsZ

A synthetic fragment encoding the promoter region of the *Sinorhizobium meliloti ftsZ* was digested by appropriate enzymes and cloned into p*lacZ290* (*Eco*RI/*Xba*I).

### Synthetic fragments used in the study

#### Ehrlichia canis P_ftsZ_

CGACTGAGACGCTCACAAagaattcaTTGCTTATATTATAACTTTCAATTCAAGTTCAATTTTACGTTGTAAAAAATCACTTTTAAATTATTAATATTAATATAAATATTATGCCATAAATTTATTATTAGTTAAATATACATACTTAATTTATTCCATATTAATCAAATCATTTGCATTATTTTATTGACGTAGTATATTAAGTTATCTTTATTATGTGTTCAGTATGTCTTTAAATATTTGTTTACCAGACCAATCCTTACTAAGACCCAGAATTACAGTTTTTGGTGTAGGTGGTGCTGGTGGTAATGCTGTAAATAACATGATACAGTCTAATCTGCACGGCGTTAACTTTGTAGTAGCTAACACAGATGCACAAGCATTAGAACTTTCCTTGTCAGAAAAGAAAATTCAGCTAGGTATCGGAatctagaaTATAGTGAGTCGTATTA AT

#### Ehrlichia canis P_ctrA_

CGACTGAGACGCTCACAAagaattcaTTTGGGGTAATATTGTGTTATATATGCAACACAAGGAAAGTATATTACATGTAATTTACTATTATACTTGAATAGCATAGTAACACTATGTATTAACTTTTGATGAAATGTTTAGTTGTTATAGGGTGATTATGTCTTTAGATTAGTGCTAAGTGCTTTATGTGTTAAGTTTGTTAATTATTGGGTTTTTAGAGGTATTTATGCGTATATTATTAATAGAAGATGATATTGCATGTGCAAAGGCAGTAGAAGCTTCTTTATCTTCAGAAGGGCATTTTTGTGAGACTATGACTTCTGCGCAGGATTGTTATGGTAGTATCATTCCGAAGAATGATGACTATGATGTAGTGATATTAGATATACATTTTCCTGGAAAAATAGATGGATATGATATTTTAGTGatctagaaTATAGTGAGTCGTAT TAAT

#### Rickettsia Prowazekii P_ftsZ_

CGACTGAGACGCTCACAAagaattcaCTATAAAAGAAAATAATCCACATTTTTCATAAAAAGTTTATGGAAATTTCTAAATCTTTTATTATACTTAAACATTAATTATATAAAATAAAAGAGTATCATGGTTTTAAATATCAAAGCTCCTGAAAATATAGTATTAAAGCCGACCATTACAGTTTTTGGCGTTGGGGGTGCAGGTAGCAATGCAGTCAATAATATGATTAGTGCTAATCTGCAAGGTGCTAATTTTGTAGTAGCTAATACTGATGCACAATCGCTTGAACATTCTTTATGCACTAACAAAATACAACTCGGTGTTTCTatctagaaTATAGTGAGTCGTATTAAT

#### Rickettsia Prowazekii P_ctrA_

CGACTGAGACGCTCACAAagaattcaAAATCTCGTTTTAAACGCCATAGTTTTTAAGTCACAGATAGAGCTTTAATAGTAAAAATTTTATCAAGATTAAATATTCTTTACTTATGTTATTAATGAATGTATAAAGCACTAGATAAAAAATTATAATTTAGTTAACAATAATTAACTGTAATTAAATTATAGCACAATATAAAAATTAAATTTTATGGAGAAAAATAATGAGAGTTTTGTTAATTGAAGACGAGCCGGAAATGGCTAACTTAATTGAGTTAACTTTAGCTTCAGAAGGGATAGTTTGTGATAAAGCTTCAGTTGGTGTGGAAGGTTTAAGACTTGTTAAAGTCGGTGGTTATGATCTAGTGATTTTAGACCTTATGTTACCGGATATTAATGGTTTTGAGATATTACTAAGATTGCGTatctagaaTATAGTGAGTCGTAT TAAT

#### Sinorhizobium meliloti Rm2011_P_ctrA_

CGACTGAGACGCTCACAAagaattcaGGCCACCAGACGTTGCTCGGCCCCGACCTGACGCCGCACGATCCGGATGCCGACCCCGAGGCGGACGAGCACCAATCGTAAAGATT

GACGCCCGCACTTAATCCCGAATTAACCTTTTGCCTTCAGATGTCGTTTGGTTAATGAATCGGGGCGAGCCGCGAGCCAGGGGTGCTGGCGGCCTCGCCAACAGGGGACGACGCGATGCGGGTGTTGTTGATCGAGGACGAGCCGACCACGGCCAAGGCGATCGAGCTGATGCTCGGCACCGAGGGGTTCAACGTCTATACGACCGACCTTGGCGAGGAGGGCTTGGACCTCGGCAAGCTGTACGACTACGACATCATCCTGCTCGACCTCAACCTGCCGGACATGCACGGCTATGACGTGCTGAAGAAGCTGCGCatctagaaTATAGTGAGTCGTATTAAT

#### Sinorhizobium meliloti Rm2011_P_ftsZ_

CGACTGAGACGCTCACAAagaattcaGTGGCACCCCAGCAAACCACCGTTCATCGCCAGCGTGGCATGGGCATGGTCCGCAAGCTGATTCAGGCGGCGCGGGCAAATTATTAAGCGCCGATGACCTTTCGCGCTGTTTTCCTATGGAAGAGCGTGAATCGTTGAGGCAAAAGGTTAATCAGGGTCCCGGCGCGCAATTCCGGACAGGGGCGGAGAAAGTTTAGGCCATGAGCATCGAATTCATCACTCCCGAGGTCGACGAACTGACCCCCCGTATCGCCGTGATCGGCGTGGGCGGCGCGGGTGGCAACGCCATCGCGAACATGATGCGTGCCGACGTGCAGGGCGTCGACTTCCTGGTCGCGAACACCGACGCGCAGGCGCTGAAGTCGTCGATCGCGCCGCAGCGCATCCAGCTGGGTGCGAAGatctagaaTATAGTGAGTCGTATTAAT

### Strain constructions

pNPTS138 derivatives were first introduced into *C. crescentus* strains by intergeneric conjugation (bi-parental mattings) and then plated on PYE harboring kanamycin (to select for recombinants) and nalidixic acid to counter select *E. coli* S17-1 donor cells^30^. A single homologous recombination event of kanamycin-resistant colonies was verified by PCR. The resulting strain was grown to stationary phase in PYE medium lacking kanamycin. Cells were then plated on PYE supplemented with 3% sucrose and incubated at 30°C. Single colonies were picked and transferred in parallel onto plain PYE plates and PYE plates containing kanamycin. Kanamycin-sensitive cells, which had lost the integrated plasmid due to a second recombination event, were then identified for disruption of the specified locus by PCR using couple of primers external to the DNA fragments used for the pNPTS138 constructs.

### Motility suppressors of NA1000 Δ*gcrB* Δ*gcrA*::Ω and mutant cells

Spontaneous mutations that suppress the motility defect of the Δ*gcrB* Δ*gcrA*::Ω mutant cell appeared as “flares” that emanated from non-motile colonies after approximately 5 to 6 days of incubation at 30°C. Three isolates (UG4211, UG4225, UG4226) were subjected to whole genome sequencing, and mutations in the *hprk* (*hprK^D47E^*), *pstS* (*pstS^G61S^*), *phoD* (*phoD^V117A^*), *phoB* (*phoB^T74A^*) and *dnaA* (dnaA^H334R^) genes were identified and confirmed by a second sanger sequencing reaction (Fasteris SA, Geneva, Switzerland). In addition, the absence of this mutation in the parental Δ*gcrB* Δ*gcrA*::Ω strain were also confirmed by a sanger sequencing reaction. Description and distribution of identified mutations are listed and described in table below.

**Table 1.**
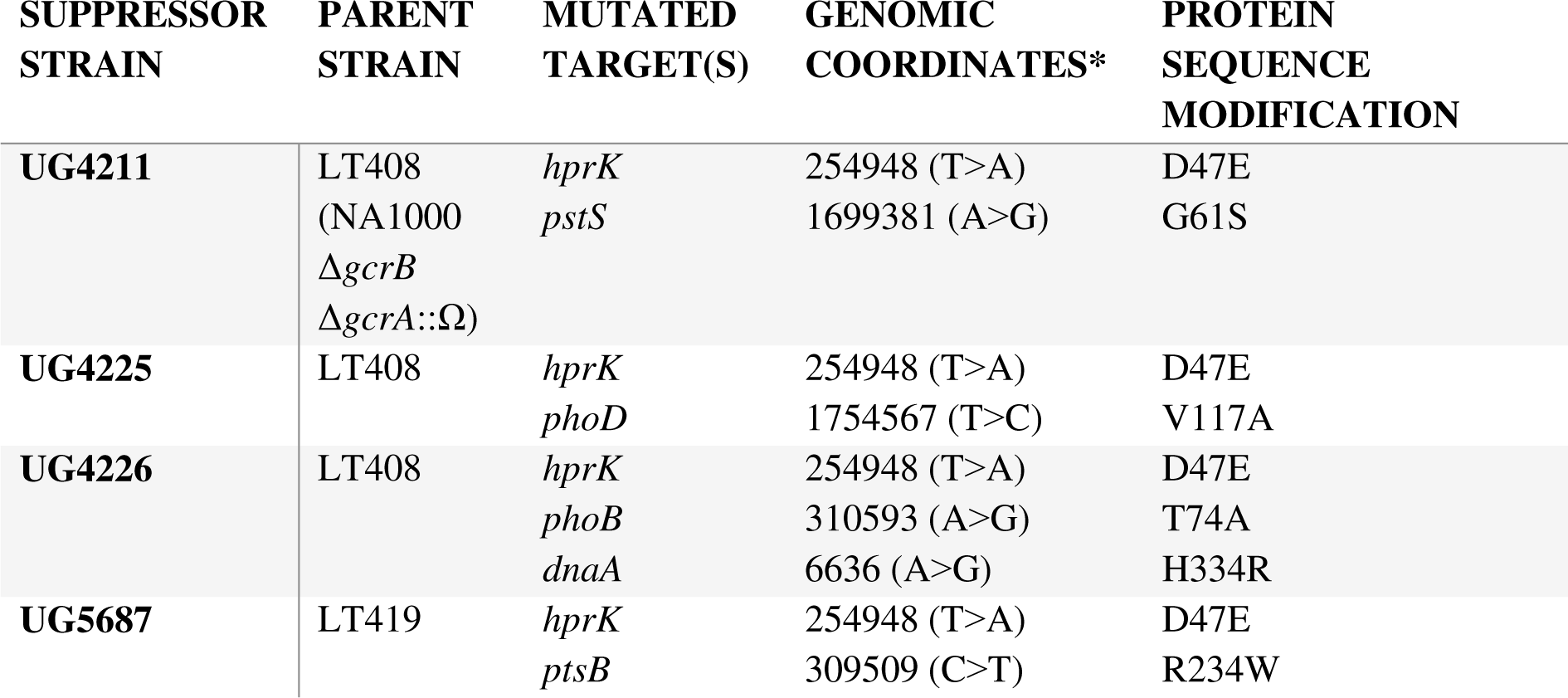
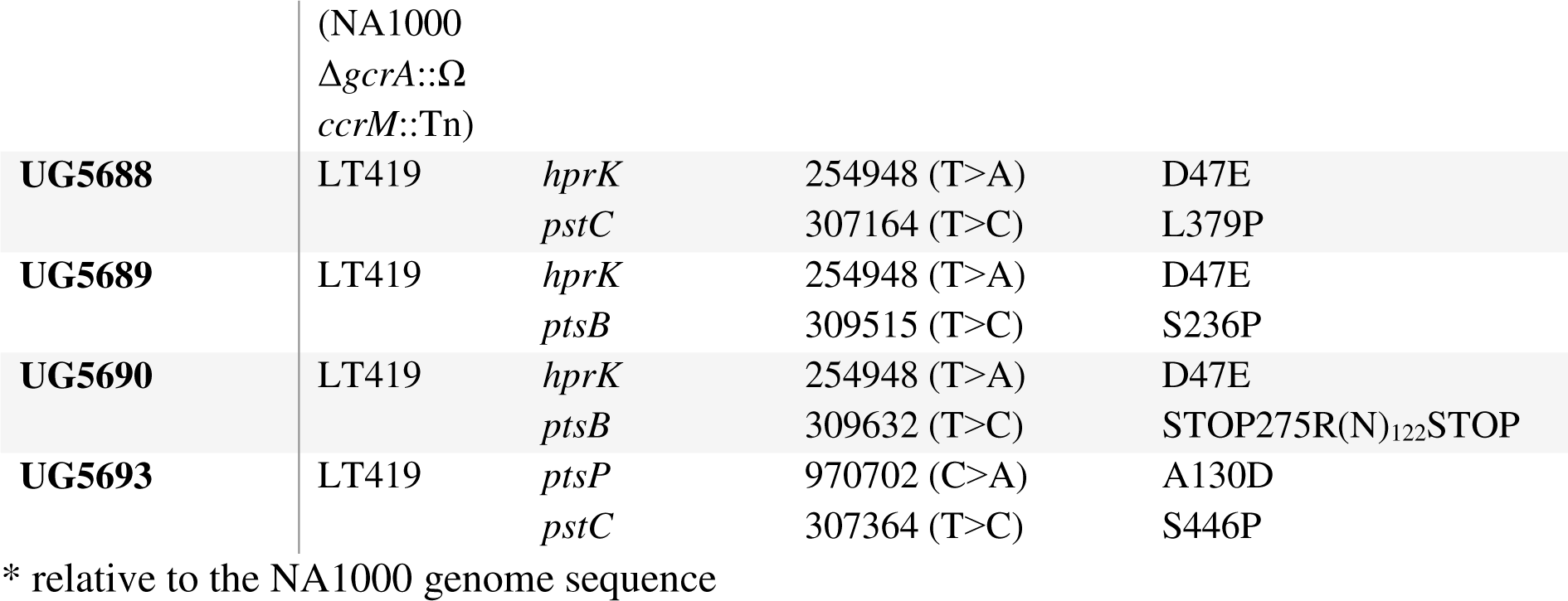
Identified mutations in the NA1000 ΔgcrB ΔgcrA::Ω and NA1000 ΔgcrA::Ω ccrM::Tn motility suppressors.

### Motility suppressors of NA1000 Δ*gcrA*::Ω *ccrM::tn* mutant cells

Spontaneous mutations that suppress the motility defect of the Δ*gcrA*::Ω *ccrM::tn* mutant cell appeared as “flares” that emanated from non-motile colonies after approximately 4 to 5 days of incubation at 30°C. Five isolates (UG5687, UG5688, UG5689, UG5690 and UG5693) were subjected to whole genome sequencing and mutations were identified in *hprk* (*hprK^D47E^*), *ptsP* (*ptsP^A130D^*), *pstB (pstB^R234W^* in UG5687, *pstB^S236P^* in UG5689 and *pstB^stop275R^* in UG5690) and *pstC* (*pstC^L379P^* in UG5688 or *pstC^S446P^* in UG5693). A second sanger sequencing reaction (Fasteris SA, Geneva, Switzerland) confirmed the presence of the identified mutations in the five suppressor isolates, and also confirmed their absence in the parental Δ*gcrA*::Ω *ccrM*::Tn strain. Description and distribution of identified mutations are listed and described in table below.

### Motility suppressors of NA1000 Δ*phoB* Δ*spoT* Δ*gcrA*::Ω mutant cells

Spontaneous mutations that suppress the motility defect of the Δ*phoB* Δ*spoT* Δ*gcrA*::Ω mutant cell appeared as “flares” that emanated from non-motile colonies after approximately 4 to 5 days of incubation at 30°C. Three isolates (MD2367, MD2369 and MD2372) were subjected to whole genome sequencing and mutations were identified in *rpoB* (*rpoB^H559Y^*) in MD2367 and MD2372 and *rpoB* (*rpoB^T561A^*) in MD2369. A second sanger sequencing reaction (Fasteris SA, Geneva, Switzerland) confirmed the presence of the identified mutations in the three suppressor isolates, and also confirmed their absence in the parental Δ*phoB* Δ*spoT* Δ*gcrA*::Ω strain using the primers rpoB-seq1 and rpoB-seq2.

### Isolation of Δ*gcrA*::Ω *hprK*^D47E^ mutant strain

Δ*gcrA*::Ω and Δ*gcrB* Δ*gcrA*::Ω overnight cultures were isolated on PYE plates, respectively, and then incubated at 30°C. The hundred clones that seemed to have a slight growth advantage (size) on plates after 3 days of incubation at 30°C were checked by PCR using hprK-seq1 and hprK-seq2 couple of primers and then submitted to sanger sequencing. A single clone with a mutation in the *hprK^D47E^* gene could be identified in Δ*gcrA*::Ω by this approach.

### β-Galactosidase assays

β-Galactosidase assays were performed at 30°C as described previously^31, 34^. Cells (50 to 200 µL) at OD_660nm_ =0.1-0.5 were lysed with chloroform and mixed with Z buffer (60 mM Na_2_HPO_4_, 40 mM NaH_2_PO_4_, 10 mM KCl and 1 mM MgSO_4_ heptahydrate) to a final volume 800 µL. 200 µL of ONPG (4 mg/mL o-nitrophenyl-β-D-galactopyranoside in 0.1 M KPO_4_ pH7.0) was added and the reaction timed. When a medium-yellow color developed, the reaction was stopped with 400 µL of 1M Na_2_CO_3_. The OD_420nm_ of the supernatant was determined and the units were calculated with the equation: U= (OD_420nm_ * 1000) / (OD_660nm_ * time (in min) * volume of culture (in mL)). Experimental values represent the averages of at least 3 independent experiments and error was computed as standard deviation (SD). p*lacZ*290 plasmids were introduced into *C. crescentus* strains by electroporation and into S. *meliloti* by bi-parental mating.

### PhoB purification and antibody production

His6-(SUMO-)PhoB protein was expressed from pCWR350 in *E. coli* Rosetta (DE3)/pLysS (Novagen, Madison, WI) and purified under native conditions using Ni2+ chelate chromatography. Briefly, cells were pelleted, resuspended in 25 mL of lysis buffer (10 mM Tris HCl pH8, 0.1 M NaCl, 1.0 mM β-mercaptoethanol, 5% glycerol, 0.5 mM imidazole triton 0.02%). Cells were sonicated (Sonifier Cell Disruptor B-30; Branson Sonic Power. Co., Danbury, CT) on ice using 12 bursts of 20 sec at output level 4.5. After centrifugation at 6’000 rpm for 20 min, the supernatant was loaded onto a column containing 5 ml of Ni-NTA agarose resin. The column was rinsed with lysis buffer, 400 mM NaCl and 10 mM imidazole, both prepared in lysis buffer. The eluted His6-(SUMO-)PhoB was used as antigen for the production of polyclonal antibodies in New Zealand white rabbits (Josman LLC, Napa, CA).

### Immunoblot analysis

Protein samples were separated by SDS-PAGE and blotted on PVDF (polyvinylidenfluoride) membranes (Merck Millipore). Membranes were blocked for 1h with PBS, 0,1% Tween 20 and 5% dry milk and then incubated for overnight with the primary antibodies diluted in TBS, 0,1% Tween 20, 5% dry milk. The different antisera were used at the following dilutions: anti-DnaA (1:20000)^35^, anti-GcrA (1:5000)^36^, anti-PhoB (1:5000), anti-MreB (1:20000)^37^, anti-MipZ (1:5000)^38^, anti-PodJ (1:10000)^39^, anti-FtsN (1:10000)^40^, anti-CtrA (1:10000)^41^, anti-CcrM (1:10000)^42^, anti-ZitP (1:10000)^43^ and anti-FljK (1:20,000)^44^. The membranes were washed 4 times for 10 min in TBS and incubated for 1h with the secondary antibody diluted in TBS, 0.1% Tween 20 and 5% dry milk. The membranes were finally washed again 4 times for 10 min in TBS and revealed with Immobilon Western Blotting Chemoluminescence HRP substrate (Merck Millipore) and Super RX-film (Fujifilm) or luminescent image analyzer (ChemidocTm MP, Biorad).

### Microscopy

PYE or PYE/5 cultivated cells in exponential growth phase were immobilized using a thin layer of 1% agarose. Phase microscopy images were taken with an Alpha Plan-Apochromatic 100X/1.46 DIC(UV) VIS-IR oil objective on an Axio Imager M2 microscope (Zeiss) and a Photometrics Evolve camera (Photometrics) controlled through Metamorph V7.5 (Universal Imaging). Images were processed using Metamorph V7.5.

### Flow cytometry analysis

PYE or PYE/5 cultivated cells in exponential growth phase (OD_660nm_ =0.3-0.6) were fixed into ice cold Ethanol solution 70% (final concentration). Fixed cells were resuspended in FACS Staining buffer pH7.2 (10 mM Tris-HCl, 1 mM EDTA, 50 mM NaCitrate, 0.01% TritonX-100) and then treated with RNase A (Roche) at 0.1 mg/mL during 30 min at room temperature. Cells were stained in FACS Staining buffer containing 0.5 µM of SYTOX Green nucleic acid stain solution (Invitrogen) and then analyzed using a BD Accuri C6 flow cytometer instrument (BD Biosciences, San Jose, California, USA). Flow cytometry data were acquired and analyzed using the CFlow Plus V1.0.264.15 software (Accuri Cytometers Inc.) and FlowJo software. 20000 cells were analyzed from each biological sample. The Green fluorescence (FL1-A) parameters were used to estimate cell chromosome contents. Relative chromosome number was directly estimated from FL1-A value of NA1000 cells treated with 20 µg/mL Rifampicin during 3 hours at 30°C. Rifampicin treatment of cells blocks the initiation of chromosomal replication but allows ongoing rounds of replication to finish.

### Motility assays and phage infectivity tests

Swarming properties were assessed with 1 μL drops of overnight culture, adjusted to an OD_660nm_ of 1, spotted on PYE soft agar plates (0.3% agar) and incubated at 30°C for 48 to 72 hours. Phage susceptibility assays were conducted by mixing 200 μL of overnight culture in 6 mL soft PYE agar and overlaid on a PYE agar plate. Upon solidification of the soft (top) agar, we spotted 4 μL drops of serial dilution of phages (ϕCbK or ϕCr30) and scored for plaques after one day incubation at 30°C.

### ϕCr30 Transduction of theΔ*gcrA* mutation

Δ*gcrA*::Ω (SpcR) transducing phage stock is a φCr30 lysate of LS3707 [24]. 300 μL of overnight cultures were infected with 30 µL of φCr30 phage stock (∼1010 pfu/ml) and incubated for 2 h at 30°C, and then plated on solid PYE containing spectinomycin/streptomycin antibiotics and 50 μM of Vanillate. Plates were incubated at 30°C, and visible colonies were counted each day. Experimental values represent the average of three biological replicates. Strains harboring the pMT335-*gcrA* complementation plasmid served as a positive control of φCr30 transduction efficiency.

### Viability test on plates

*WT* and mutants were freshly electroporated with pMT335, pMT335-*gcrA*, pMT335-*ctrA* and pMT335-*ctrA*Δ3Ω plasmids, and then plated on solid PYE containing 1 μg/mL of gentamycin. Ten colonies are then collected for each condition, then mixed and resuspended in 50 µL liquid PYE. 4 μL drops of serial dilutions (10^-1^ to 10^-6^) of liquid culture adjusted to OD_660nm_ =0.1 (NanoDrop, cell culture measurement) are then spotted on PYE plates supplemented with 1 μg/mL of gentamycin with or without 50 µM vanillate.

### Shearing experiments

PYE overnight culture of *C. crescentus* NA1000 *WT* and derivatives were harvested and washed 3 times with PYE, and then freshly restarted in PYE or PYE/5 for 5 h at 30°C. The equivalent of 5 mL of exponential phase cultures at 0.5 OD_660nm_ of *WT* or mutant strains were harvested and resuspended in 1ml of PYE. Then, we pumped in and out (10x) the cells into a syringe endowed with a thin diameter needle. We centrifuged the shear-stressed cells to remove cells debris and collected 200 μL of each supernatant. We added SDS-blue straining and loaded samples on SDS PAGE gels.

### Chromatin ImmunoPrecipitation coupled to deep Sequencing (ChIP-Seq)

Cultures of exponentially growing (OD660nm of 0.5) *C. crescentus* NA1000 *WT* and *ΔpstS* strains were treated with formaldehyde (1% final concentration) in 10 μM sodium phosphate buffer (pH 7.6) at RT for 10 min to achieve crosslinking. Subsequently, the cultures were incubated for an additional 30 min on ice and washed three times in phosphate buffered saline (PBS, pH 7.4). The resulting cell pellets were stored at -80°C. After resuspension of the cells in TES buffer (10 mM Tris-HCl pH 7.5, 1 mM EDTA, 100 mM NaCl) containing 10 mM of DTT, they were incubated in the presence of Ready-Lyse lysozyme solution (Epicentre, Madison, WI) for 10 minutes at 37°C, according to the manufacturer’s instructions. Lysates were sonicated (Bioruptor® Pico) at 4°C using 15 bursts of 30 sec to shear DNA fragments to an average length of 0.3-0.5 kbp and cleared by centrifugation at 14,000 rpm for 2 min at 4°C. The volume of the lysates was then adjusted (relative to the protein concentration) to 1 ml using ChIP buffer (0.01% SDS, 1.1% Triton X-84 100, 1.2 mM EDTA, 16.7 mM Tris-HCl [pH 8.1], 167 mM NaCl) containing protease inhibitors (Roche) and pre-cleared with 80 μl of Protein-A agarose (Roche, www.roche.com) and 100 μg BSA. 5% of each pre-cleared lysate was kept as total input samples (negative control samples). The rest of the pre-cleared lysates was incubated overnight at 4°C with polyclonal rabbit antibodies targeting PhoB (1:1,000 dilution). The immuno-complexes were captured after incubation with Protein-A agarose (pre-saturated with BSA) during a 2 h incubation at 4°C and they were washed subsequently with low salt washing buffer (0.1% SDS, 1% Triton X-100, 2 mM EDTA, 20 mM Tris-HCl pH 8.1, 150 mM NaCl), with high salt washing buffer (0.1% SDS, 1% Triton X-100, 2 mM EDTA, 20 mM Tris-HCl pH 8.1, 500 mM NaCl), with LiCl washing buffer (0.25 M LiCl, 1% NP-40, 1% deoxycholate, 1 mM EDTA, 10 mM Tris-HCl pH 8.1) and finally twice with TE buffer (10 mM Tris-HCl pH 8.1, 1 mM EDTA). The complexes were eluted from the Protein-A agarose beads with two times 250 μl elution buffer (SDS 1%, 0.1 M NaHCO3, freshly prepared) and then, just like total input samples, incubated overnight with 300 mM NaCl at 65°C to reverse the crosslinks. The samples were then treated with 2 μg of Proteinase K for 2 h at 45°C in 40 mM EDTA and 40 mM Tris-HCl (pH 6.5). DNA was extracted using phenol:chloroform:isoamyl alcohol (25:24:1), ethanol-precipitated using 20 μg of glycogen as a carrier and resuspended in 50 μl of DNAse/RNAse free water.

Immunoprecipitated chromatin was used to prepare sample libraries used for deep-sequencing at Fasteris SA (Geneva, Switzerland). ChIP-Seq libraries were prepared using the DNA Sample Prep Kit (Illumina) according to the manufacturer’s instructions. Single-end runs (50 cycles) were performed on an Illumina Genome Analyzer IIx or HiSeq2000, yielding several million reads. The single-end sequence reads (stored as FastQ files) were mapped against the genome of *C. crescentus* NA1000 (NC_011916.1) using Bowtie version 0.12.9 (-qS -m 1 parameters, http://bowtie-bio.sourceforge.net/). ChIPseq read sequencing and alignment statistics are summarized in Suppl. Table 1.

The standard genomic position format files (BAM, using Samtools, http://samtools.sourceforge.net/) were imported into SeqMonk version 0.34.1 (Braham http://www.bioinformatics.babraham.ac.uk/projects/seqmonk/) to build ChIPseq normalized sequence read profiles. Briefly, the genome was subdivided into 50 bp probes, and for every probe, we calculated the number of reads per probe as a function of the total number of reads (per million, using the Read Count Quantitation option). Analyzed data as shown in Figure S5 are provided in Suppl. Table 2.

Using the web-based analysis platform Galaxy (https://usegalaxy.org), PhoB ChIP-Seq peaks were called using MACS2^45^ relative to the total input DNA samples. The q-value (false discovery rate, FDR) cut-off for called peaks was 0.05. Peaks were rank-ordered according to fold-enrichment (Suppl. Table 3) and peaks with a fold-enrichment values >5 were retained for further analysis. >5 factor was selected as it allows to retain highly statistical peaks in both WT and Δ*pstS* datasets. In addition, this >5 factor selection allows to enrich, in the PhoB ChIP-seq list, specific loci presenting a >2-fold enrichment between WT and Δ*pstS* conditions. Consensus sequences common to the 98 enriched PhoB-associated loci (Δ*pstS* MACS2 datasets) were identified by scanning peak sequences (+ or - 75 bp relative to the coordinates of the peak summit) for conserved motifs using MEME^22^ (MEME-ChIP, Distribution mode : Any Number of Repetition). Sequence data have been deposited to the Gene Expression Omnibus (GEO) database.

### Chromatin ImmunoPrecipitation followed by quantitative PCR

Chromatin ImmunoPrecipitation of *C. crescentus* NA1000 *WT*, UG4211, UG4226, MD138 and MD435 were prepared as described above. Pre-clearing and immune-complexes capture were carried out, depending on the origin of the antibody used, with Protein-A (Rabbit antibodies) or Protein-G (Mouse antibodies) agarose beads, respectively. GcrA (1:1,000 dilution, rabbit)^36^and RpoD (1:250 dilution, Mouse IgG2b) (Neoclone, BioLegend) antibodies were used. Real-time PCR was performed using the Step-One Real-Time PCR system (Applied Biosystems) using ChIP sample at different DNA dilutions (5 µL), with 12.5 µL of SYBR green PCR master mix (Quanta Biosciences), 0.5 µL of each primer (10 µM), and 6.5 µL of water per reaction. PCR assay parameters were one cycle at 95°C for 5 min followed by 40 cycles at 95°C for 15 s, 55°C for 20 s, and 60°C for 20 s. Serial dilutions of chromatin total-input samples were used to generate the standard curve. GcrA and RpoD (σ^70^) occupancy at the *ftsN* and *pstC* promoter regions were normalized (ratio) by the *ter* locus quantification (DNA input normalization). Average values are from triplicate measurements from two independent biological replicates. The DNA regions analyzed by real-time PCR were from nucleotide −75 to +4 relative to the start codon of *ftsN*, from −131 to +122 relative to the start codon of *pstC* and from position 1971139 to 1971313 on the *C. crescentus* NA1000 chromosome for the terminus region (*ter*).

### Transposon suppressor screen coupled to deep Sequencing (Tn-Seq)

Transposon mutagenesis of *ΔccrM*::Ω strain was done by intergeneric conjugation from *E. coli* S17-1 *λ*pir harbouring the himar1-derivative pHPV414^46^. A Tn-library of >20,000 kanamycin- and nalidixic-acid-resistant clones was collected. Clones were harvested and chromosomal DNA was extracted. Genomic DNA was used to generate barcoded Tn-Seq libraries and submitted to Illumina HiSeq 2000 sequencing (Fasteris SA). Tn-insertion specific reads (50 bp long) were sequenced using the himar-Tnseq primer (5’-AGACCGGGGACTTATCAGCCAACCTGTTA-3’). The million reads generated by sequencing were mapped using the web-based analysis platform Galaxy (https://usegalaxy.org; Map_with_Bowtie_for_Illumina_V1.1.2) to the *C. crescentus* NA1000 genome (NC_011916.1). Using Samtool_V0.1.18, BED file format encompassing the Tn insertion coordinates were generated and then imported into SeqMonk V1.40.0 (www.bioinformatics.babraham.ac.uk/projects/) to assess the total number of Tn insertion per chromosome position (Tn-insertion per millions of reads count) or per coding sequence (CDS). For CDS Tn-insertion ratio calculation, SeqMonk datasets were exported into Microsoft Excel files (Dataset S1) for further analyses, as described previously^7^. Briefly, to circumvent ratio issue for a CDS Tn-insertion value of 0 and CDS that do not share sufficient statistical Tn-insertions, an average value of all CDS-Tn insertions normalized to the gene size was calculated, and 1% of this normalized value was used to correct each CDS-Tn insertion value.

### Cell generation time determination

Cells growth in PYE medium was done in an incubator at 30°C under agitation (190 rpm) and monitored at OD_660nm_. Generation time values were extracted from the curves using the Doubling Time application (http://www.doubling-time.com). Values represent the averages of at least 3 independent clones.

**Table 2.** Plasmids used in this study.

| Plasmid | Relevant characteristics | Reference or source |
| --- | --- | --- |
| pNPTS138 | Suicide vector used for gene replacement; Kan<sup>R</sup> | M.R.K. Alley unpublished |
| pNPTS138_Δ*phoB* | pNPTS138 derivative carrying the in-frame deletion of *CCNA_00296*; Kan<sup>R</sup> | This work |
| pNPTS138_Δ*pstS* | pNPTS138 derivative carrying the in-frame deletion of *CCNA_01583*; Kan<sup>R</sup> | This work |
| pNPTS138_Δ*phoD* | pNPTS138 derivative carrying the in-frame deletion of *CCNA_00164*; Kan<sup>R</sup> | This work |
| pNPTS138-spoT-us | pNPTS138 derivative carrying the in-frame deletion of *pkk2*; Kan<sup>R</sup> | This work |
| pMT384 | Integrative plasmid at the *vanA* locus; Kan<sup>R</sup> | 47 |
| pMT384-*pstS* | pMT384 derivative carrying the *pstS* ORF, Kan<sup>R</sup> | This work |
| pMT384-*pstS*<sup>G61S</sup> | pMT384 derivative carrying the *pstS* G61S variant, Kan<sup>R</sup> | This work |
| pMT335 | Medium copy number plasmid for inducible expression; P<sub>van</sub>, Gm<sup>R</sup> | 47 |
| pMT335-*hprK* | pMT335 derivative carrying the *hprK* ORF; Gm<sup>R</sup> | This work |
| pMT335-*hprK*<sup>D47E</sup> | pMT335 derivative carrying the *hprK* D47E variant; Gm<sup>R</sup> | This work |
| pMT335-*hprK1.4* | pMT335 derivative carrying the *hprK1.4* variant; Gm<sup>R</sup> | This work |
| pMT335-*pstS* | pMT335 derivative carrying the *pstS* ORF, Gm<sup>R</sup> | This work |
| pMT335-*pstS*<sup>G61S</sup> | pMT335 derivative carrying the *pstS* G61S variant, Gm<sup>R</sup> | This work |
| pMT335-*gcrA* | pMT335 derivative carrying the *gcrA* ORF; Gm<sup>R</sup> | 7 |
| pMT335-*ctrA* | pMT335 derivative carrying the *ctrA* ORF; Gm<sup>R</sup> | This work |
| pMT335-*ctrA*Δ3Ω | pMT335 derivative carrying the non-degradable form of *ctrA*; Gm<sup>R</sup> | This work |
| PlacZ290 | Low copy number vector containing *lacZ* gene (used to create *lacZ* transcriptional fusion); Tet<sup>R</sup> | 48 |
| PlacZ290-P<sub>mipZ</sub> | PlacZ290 containing *mipZ* promoter region | 6 |
| PlacZ290-P<sub>podJ</sub> | PlacZ290 containing *podJ* promoter region | 6 |
| PlacZ290-P<sub>pleC</sub> | PlacZ290 containing *pleC* promoter region | 6 |
| PlacZ290-P<sub>1149</sub> | PlacZ290 containing *CCNA_01149* promoter region | 49 |
| PlacZ290-P<sub>pftsZ</sub> | PlacZ290 containing *ftsZ* promoter region | 50 |
| PlacZ290-P<sub>tipF</sub> | PlacZ290 containing *tipF* promoter region | 6 |
| *PlacZ290-P<sub>pctrA</sub>* | *PlacZ290* containing *ctrA* promoter region | 51 |
| *PlacZ290-P<sub>EcctrA</sub>* | *PlacZ290* containing *Ehrlichia canis ctrA* promoter | This work |
| *PlacZ290-P<sub>EcftsZ</sub>* | *PlacZ290* containing *Ehrlichia canis ftsZ* promoter | This work |
| *PlacZ290-P<sub>RpctrA</sub>* | *PlacZ290* containing *Rickettsia Prowazekii ctrA* promoter region | This work |
| *PlacZ290-P<sub>RpftsZ</sub>* | *PlacZ290* containing *Rickettsia Prowazekii ftsZ* promoter region | This work |
| *PlacZ290-P<sub>Rm2011ctrA</sub>* | *PlacZ290* containing *Sinorhizobium meliloti ctrA* promoter region | This work |
| *PlacZ290-P<sub>Rm2011ftsZ</sub>* | *PlacZ290* containing *Sinorhizobium meliloti ftsZ* promoter region | This work |
| *PlacZ290-P<sub>pstC</sub>* | *PlacZ290* containing *pstC* promoter region (-454 to +123 relative to the ATG) | This work |
| *PlacZ290-P<sub>rpoD</sub>* | *PlacZ290* containing *rpoD* promoter region (-695 to +28 relative to the ATG) | This work |
| *PlacZ290-P<sub>ftsN</sub>* | *PlacZ290* containing *ftsN* promoter region | 7 |
| *PlacZ290-P<sub>ftsN\*</sub>* | *PlacZ290-pftsN* harboring the A→T mutation in position -52 relative to the ATG (GATTC→GTTTC) | 7 |
| pXTCYC4 | Non replicative vector in *C. crescentus* for expression of genes under the control of Pxyl after the integration at chromosomal *xylX* locus; Gm<sup>R</sup> | 47 |
| pXTCYC4-*relA*'-FLAG | pXTCYC4 containing *relA*'-FLAG | 52 |

**Table 3.** Strains used in this study.

| Strains | Relevant characteristics | Reference or source |
| --- | --- | --- |
| *Escherichia coli* |  |  |
| S17 | RP4, Tc::Mu Km::Tn7 | 30 |
| EC100D | *F- mcrA Δ(mrr-hsdRMS-mcrBC) Φ80dlacZΔM15 ΔlacX74 recA1 endA1 araD139 Δ(ara, leu)7697 galU galK λ- rpsL (StrR) nupG* | Epicentre |
| *Caulobacter crescentus* |  |  |
| NA1000 | Synchronizable variant of strain CB15 | 29 |
| LS3707 | NA1000 $\Delta gcrA::\Omega$ *xylX::Pxyl-gcrA* | 36 |
| LT406 | NA1000 $\Delta gcrB$ | 7 |
| LT407 | NA1000 $\Delta gcrA::\Omega$ | 7 |
| LT408 | NA1000 $\Delta gcrB \Delta gcrA::\Omega$ | 7 |
| UG4211 | NA1000 $\Delta gcrB \Delta gcrA::\Omega$ *hprK*<sup>D47E</sup> *pstS*<sup>G61S</sup> | This work |
| UG4225 | NA1000 $\Delta gcrB \Delta gcrA::\Omega$ *hprK*<sup>D47E</sup> *phoD*<sup>V117A</sup> | This work |
| UG4226 | NA1000 $\Delta gcrB \Delta gcrA::\Omega$ *hprK*<sup>D47E</sup> *dnaA*<sup>H334R</sup> *phoB*<sup>T74A</sup> | This work |
| GP29-73 | NA1000 $\Delta gcrA::\Omega$ *hprK*<sup>D47E</sup> | This work |
| MD138 | NA1000 $\Delta phoB$ | This work |
| MD435 | NA1000 $\Delta pstS$ | This work |
| MD516 | NA1000 $\Delta phoD$ | This work |
| GP27-21 | NA1000 *vanA::Pvan* (empty plasmid) | This work |
| GP21-48 | NA1000 *vanA::Pvan-pstS* | This work |
| GP21-50 | NA1000 *vanA::Pvan-pstS*<sup>G61S</sup> | This work |
| GP21-54 | NA1000 $\Delta pstS$ *vanA::Pvan-pstS* | This work |
| GP21-56 | NA1000 $\Delta pstS$ *vanA::Pvan-pstS*<sup>G61S</sup> | This work |
| MD454 | NA1000 $\Delta pstS \Delta gcrA::\Omega$ | This work |
| MD414 | NA1000 $\Delta phoB \Delta gcrA::\Omega$ | This work |
| MD569 | NA1000 $\Delta phoD \Delta gcrA::\Omega$ | This work |
| GP22-40 | NA1000 $\Delta hprK$ | This work |
| JC1323 | NA1000 *hprK1.4* | 9 |
| GP25-87 | NA1000 $\Delta hprK \Delta gcrA::\Omega$ | This work |
| GP25-71 | NA1000 *hprK1.4* $\Delta gcrA::\Omega$ | This work |
| MD214 | NA1000 $\Delta phoB \Delta gcrA::\Omega$ *xylX::Pxyl-gcrA* | This work |
| UG2212 | NA1000 *ccrM::Ω* | 6 |
| LT420 | NA1000 *ccrM::Tn* (399278<sup>+</sup>) | 7 |
| LT419 | NA1000 $\Delta gcrA::\Omega$ *ccrM::Tn* (399278<sup>+</sup>) | 7 |
| FC769 | NA1000 $\Delta spoT$ | 53 |
| SS3 | NA1000 $\Delta ptsP$ | 54 |
| LT1804 | NA1000 $\Delta ptsP \Delta spoT$ | Delaby *et al*,<br>submitted |
| LT1805 | NA1000 $rpoB^{H559Y}$ | Delaby *et al*,<br>submitted |
| LT1806 | NA1000 $\Delta ptsP \Delta spoT rpoB^{H559Y}$ | Delaby *et al*,<br>submitted |
| MD1158 | NA1000 $\Delta phoB \Delta spoT$ | This work |
| MD1735 | NA1000 $\Delta phoB \Delta spoT \Delta gcrA::\Omega$ | This work |
| MD2367 | NA1000 $\Delta phoB \Delta spoT \Delta gcrA::\Omega rpoB^{H559Y}$ | This work |
| MD2372 | NA1000 $\Delta phoB \Delta spoT \Delta gcrA::\Omega rpoB^{H559R}$ | This work |
| MD2369 | NA1000 $\Delta phoB \Delta spoT \Delta gcrA::\Omega rpoB^{T561A}$ | This work |
| UG5687 | NA1000 $\Delta gcrA::\Omega ccrM::tn hprK^{D47E} pstB^{R234W}$ | This work |
| UG5688 | NA1000 $\Delta gcrA::\Omega ccrM::tn hprK^{D47E} pstC^{L379P}$ | This work |
| UG5689 | NA1000 $\Delta gcrA::\Omega ccrM::tn hprK^{D47E} pstB^{S236P}$ | This work |
| UG5690 | NA1000 $\Delta gcrA::\Omega ccrM::tn hprK^{D47E} pstB^{stop275R}$ | This work |
| UG5693 | NA1000 $\Delta gcrA::\Omega ccrM::tn ptsP^{A130D} pstC^{S446P}$ | This work |
| UG1573 | NA1000 $\Delta phoB::tet$ | 55 |
| MD1126 | NA1000 $\Delta phoB::tet \Delta gcrA::\Omega$ | This work |
| GP26-17 | NA1000 $\Delta hprK \Delta phoB::tet$ | This work |
| GP26-21 | NA1000 $hprK1.4 \Delta phoB::tet$ | This work |
| GP26-33 | NA1000 $\Delta hprK \Delta phoB::tet \Delta gcrA::\Omega$ | This work |
| GP26-35 | NA1000 $hprK1.4 \Delta gcrA::\Omega \Delta phoB::tet$ | This work |
| YB767 | NA1000 $pstS::tn$ | 20 |
| MD1128 | NA1000 $pstS::tn \Delta gcrA::\Omega$ | This work |
| GP2678 | NA1000 $\Delta hprK pstS::tn$ | This work |
| GP26-87 | NA1000 $\Delta hprK \Delta gcrA::\Omega pstS::tn$ | This work |
| TPA2357 | NA1000 $\Delta fljx6$ | 56 |
| JC835 | NA1000 $xylX::pXTCYC4$ | 52 |
| JC820 | NA1000 $xylX::pXTCYC4-relA'-M2$ | 52 |
| MD2401 | NA1000 $xylX::pXTCYC4-relA'-M2 \Delta gcrA::\Omega$ | This work |
| MD2402 | NA1000 $xylX::pXTCYC4 \Delta gcrA::\Omega$ | This work |
| RH1476 | NA1000 *spoT*<sub>ΔACT</sub> | 15 |
| RH1752 | NA1000 *spoT*<sub>D81G</sub> | 14 |
| RH1844 | NA1000 *spoT*<sub>Y323A</sub> | 14 |
| RH1586 | NA1000 *spoT*<sub>Y323AΔACT</sub> | 15 |
| RH2193 | NA1000 *spoT*<sub>D81GY323A</sub> | 14 |
| MD2403 | NA1000 *rpoB*<sup>H559Y</sup> Δ*gcrA*::Ω | This work |
| MD2404 | NA1000 *rpoB*<sup>H559Y</sup> *pstS* ::*tn* | This work |
| MD2404 | NA1000 *rpoB*<sup>H559Y</sup> *pstS* ::*tn* Δ*gcrA*::Ω | This work |
| MD2405 | NA1000 *rpoB*<sup>H559Y</sup> Δ*gcrA*::Ω *pstS* ::*tn* | This work |
| MD2406 | NA1000 Δ*phoB* *xylX*::pXTCYC4 | This work |
| MD2407 | NA1000 Δ*phoB* *xylX*::pXTCYC4-*relA*'-M2 | This work |
| MD2408 | NA1000 Δ*phoB* *xylX*::pXTCYC4 Δ*gcrA*::Ω | This work |
| MD2409 | NA1000 Δ*phoB* *xylX*::pXTCYC4-*relA*'-M2 Δ*gcrA*::Ω | This work |
| MD2410 | NA1000 Δ*pstS* *xylX*::pXTCYC4 | This work |
| MD2411 | NA1000 Δ*pstS* *xylX*::pXTCYC4-*relA*'-M2 | This work |
| MD2412 | NA1000 Δ*pstS* *xylX*::pXTCYC4 Δ*gcrA*::Ω | This work |
| MD2413 | NA1000 Δ*pstS* *xylX*::pXTCYC4-*relA*'-M2 Δ*gcrA*::Ω | This work |
| MD2414 | NA1000 Δ*phoD* *xylX*::pXTCYC4 | This work |
| MD2415 | NA1000 Δ*phoD* *xylX*::pXTCYC4-*relA*'-M2 | This work |
| MD2416 | NA1000 Δ*phoD* *xylX*::pXTCYC4 Δ*gcrA*::Ω | This work |
| MD2417 | NA1000 Δ*phoD* *xylX*::pXTCYC4-*relA*'-M2 Δ*gcrA*::Ω | This work |
| *Sinorhizobium meliloti* |  |  |
| LT1387 | Rm2011 | 33 |

**Table 4.**
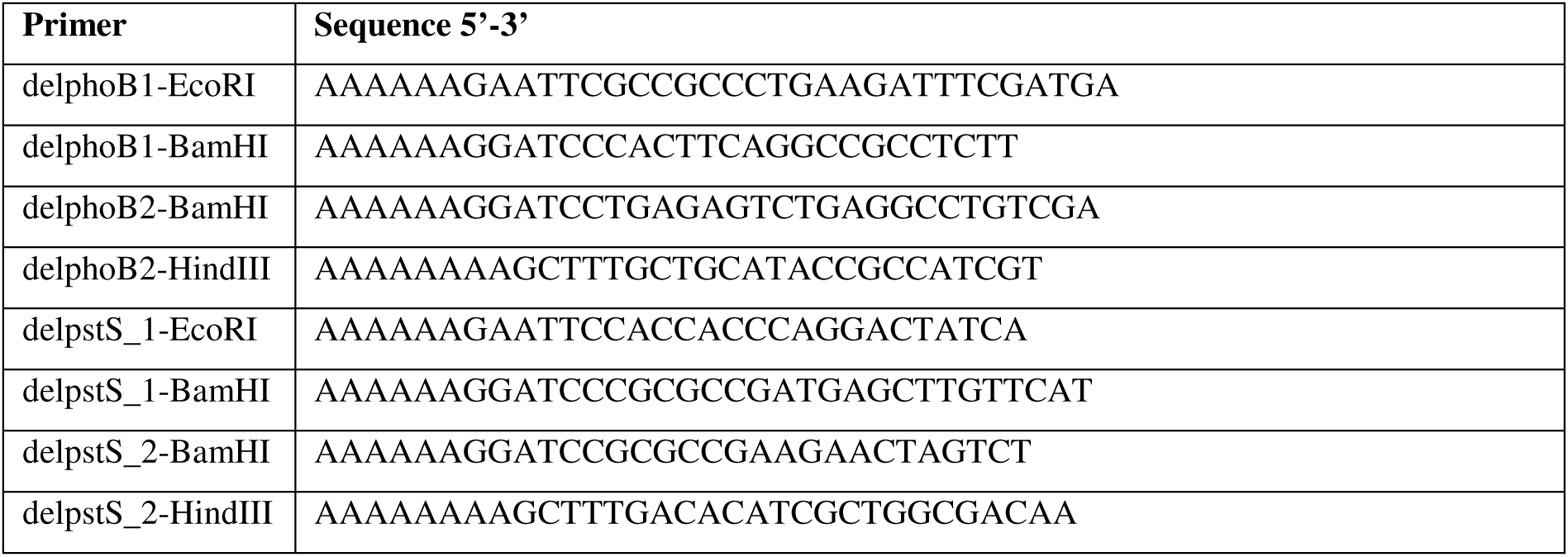
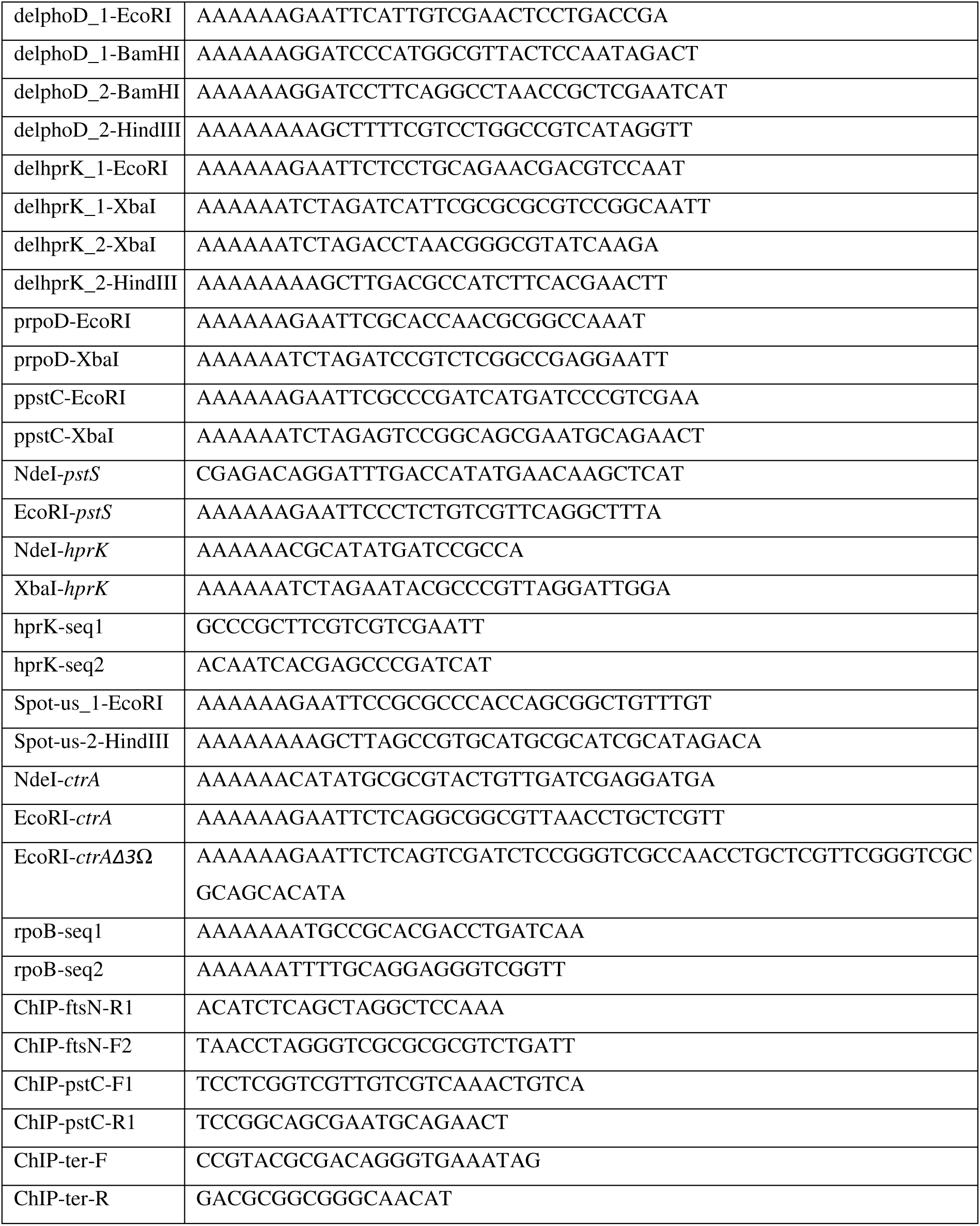
Oligonucleotides used in this study.

## Acknowledgements

Funding support is from Swiss National Science Foundation grant 31003A_182576 (to P.H.V.). We thank Regis Hallez (University of Namur, Belgium) and Justine Collier (University of Lausanne, Switzerland) for strains. We also thank Martin Thanbichler (Universty of Marburg, Germany) for providing anti-FtsN antiserum.

## Author Contributions

M.D., G.P., and P.H.V. conceived and designed the experiments. M.D., G.P., C.F. and L.D. performed the experiments. M.D., G.P., and P.H.V. analyzed the data. M.D., G.P., and P.H.V. wrote the paper.

