## Supplemental Figures for "Bypass of an epigenetic S-phase transcriptional module by convergent nutrient stress signals"

### Supplemental Figure legends

**Supplemental Figure 1. Characterization of Δ*gcrA* motility suppressors.**

(**A**) DNA content analysis as determined by FACS (FL1-A channel) of *WT*, Δ*gcrA::*Ω, Δ*gcrA::*Ω Δ*gcrB*, Δ*gcrA::*Ω Δ*gcrB hprK* pstS**, Δ*gcrA::*Ω Δ*gcrB hprK* phoD** and Δ*gcrA::*Ω Δ*gcrB hprK* phoB*dnaA** cells during exponential growth in PYE. Percentages of cells containing >2 chromosomes are also indicated. (**B**) Immunoblots showing steady-state levels of ZitP in extracts of cells grown exponentially in PYE. MreB serves as a loading control. (**C**) Sensitivity of *WT* and mutant cells to the pilus-specific phage ϕCbK and the S-layer specific phage ϕCr30. Serial dilutions of ϕCbK and ϕCr30 were spotted on lawns of cells embedded in PYE top-agar on PYE plates. The pictures on the right show the buoyancy of the cells after Percoll density gradient centrifugation. (**D**) Growth curve measurement of the three Δ*gcrA::*Ω Δ*gcrB* motility suppressors grown in PYE. Error bars are standard deviation from three biological replicates.

**Supplemental Figure 2. Simultaneous mutation of the PTS and Pho systems permits cellular cycling in the absence of GcrA and CcrM.**

(**A**) Immunoblots showing steady-state levels of PodJ, CtrA and GcrA in synchronized *WT* and Δ*gcrB* Δ*gcrA::Ω hprK* pstS** populations during exponential growth in PYE. (**B**) DNA content analysis as determined by FACS (FL1-A channel) of synchronized *WT*, Δ*gcrA::*Ω Δ*gcrB hprK* pstS**, Δ*gcrA::*Ω Δ*gcrB hprK* dnaA* phoB**, Δ*gcrA::*Ω Δ*ccrM::*Tn *hprK* pstB* and Δ*gcrA::*Ω Δ*ccrM::*Tn *ptsP* pstC** cells grown exponentially in PYE. The relative amount of chromosome (n) is indicated on the x-axis. The pictures to the right show the buoyancy of the cells after Percoll density gradient centrifugation.

**Supplemental Figure 3. Tn-insertions in *spoT* and the genes encoding the PTS system improve growth of the Δ*ccrM::* Ω cells.**

(**A**) Table summarizing the selected top targets promoting the growth of Δ*ccrM::*Ω cells compare to the *WT* as determined by Tn-Seq analysis. Ranking and relative CDS Tn-insertions enrichment (blue gradient) in Δ*ccrM::Ω vs WT* are indicated. The complete Tn-Seq analysis is available in Table S1. (**B**) Tn-insertion bias in coding sequences (CDS) of Δ*ccrM::*Ω cells relative to *WT* cells as determined by Tn-Seq. X-axis shows chromosome position and y-axis gives CDS Tn-insertion ratios. The position of selected top targets in (A) are indicated. (**C**)

Local (base pair resolution) of the Δ*ccrM::*Ω */WT* Tn-insertion bias in the coding sequences of *ftsZ*, *spoT*, *hrpK*, *CCNA_00240* (encoding PTS system component IIA), *ccrM* and *gcrA* loci are depicted. The known SpoT subdomains are shown. The red rectangle in *ccrM* delimits the area coding for the CcrM DNA methylase domain (grey) that is deleted in the Δ*ccrM::*Ω allele.

**Supplemental Figure 4. The (p)ppGpp alarmone synthesized by SpoT in PYE/5 induces the transcription of genes controlled by the GcrA/CcrM module.**

(**A**) β-galactosidase activities of $P_{ftsN}$- and $P_{ftsN*}$-*lacZ* promoter probe plasmids in *WT* and Δ*spoT* during exponential phase growth in PYE or 5 hours shifting to PYE/5. (B) β-galactosidase activities of $P_{mipZ}$-, $P_{ftsN}$- and $P_{pstC}$-*lacZ* promoter probe plasmids in exponential *WT* and Δ*pstS* cells cultivated in PYE. (**A** and **B**) β-galactosidase activities are expressed as percentage relative to *WT* measured in Miller units. Data are from four independent experiments; error bars are standard deviation.

**Supplemental Figure 5. Overexpression of GcrA is deleterious when phosphate starvation is triggered.**

(**A** and **B**) Efficiency of plating assay of *WT* and mutant cells expressing *gcrA* from the vanillate-inducible $P_{van}$ promoter on the pMT335 plasmid. (**C**) Efficiency of plating assay of *WT* and mutants expressing *ctrA* or *ctrA*Δ3Ω (non-degradable form of CtrA) from the vanillate-inducible $P_{van}$ promoter on the pMT335 plasmid**.** (-) Corresponds to the empty vector and serves as a control. Serial ten-fold dilutions were plated on PYE agar supplemented with gentamycin. When indicated, vanillate 50 µM is added.

**Supplementary Figure 6. The Δ*hprK* and *hprK1.4* mutations are not equivalent, yet they both favor growth of cells lacking GcrA.**

(**A**) Growth curve analysis of the *WT*, Δ*hprK* and *hprK1.4* strains complemented or not with the wild-type or *hprK** and *hprK1.4* point mutants (expressed from the vanillate-inducible $P_{van}$ promoter on the pMT335 plasmid) in PYE supplemented with 50 µM vanillate. Δ*hprK* and *hprK1.4* show a growth defect that is partially complemented with overexpression of *WT* HprK. (**B**) DNA content analysis as determined by FACS (FL1-A channel) of *WT*, Δ*gcrA::*Ω, Δ*gcrA::*Ω Δ*gcrB*, Δ*hprK*, *hprk1.4*, Δ*gcrA::*Ω Δ*hprK*, Δ*gcrA::*Ω *hprK1.4* , Δ*gcrA::*Ω Δ*gcrB hprK* pstS** and Δ*gcrA::*Ω Δ*gcrB hprK* dnaA* phoB** cells during exponential growth in PYE. Percentages of cells containing >2 chromosomes are also indicated. Despite an increase in G1 phase population of Δ*hprK* and *hprK1.4* cultures, Δ*hprK* Δ*gcrA::*Ω single and Δ*gcrA::*Ω *hprK1.4* double mutant cells still show an accumulation of extra-chromosomes as Δ*gcrA::*Ω cells. (**C**) β-galactosidase activities of $P_{ftsN}$- and $P_{pstC}$-*lacZ* promoter probe plasmids in *WT*, Δ*hprK* and *hprK1.4* cells cultivated in exponential phase in PYE. β-galactosidase activities are expressed as percentage relative to *WT* measured in Miller units. Data are from four independent experiments; error bars are standard deviation.

**Supplemental Figure 7. Genome-wide promoter occupancy of PhoB in phosphate-limited conditions.**

(**A**) Genome-wide occupancy of PhoB on the *C. crescentus* chromosome in *WT* and Δ*pstS* cells as determined by ChIP-Seq analysis using polyclonal antibodies to PhoB. Relevant peak numbers showing a five-fold enrichment of PhoB occupancy compared to the input DNA control are indicated. The complete PhoB ChIP-Seq analysis is available in Table S2. (**B**) Venn diagram showing a total of 121 PhoB statistically significant peaks overlapping with promoter intergenic regions, based on a five-fold enrichment of PhoB occupancy in either *WT* and/or Δ*pstS* cells *versus* their respective input DNA controls, (**C**) Venn diagram comparing the 121 selected PhoB peaks with the one published by Lubin *et al.* where an epitope-tagged PhoB-3xFLAG expressed protein and an anti-M2 antibody were used for ChIP-Seq. (**D**) PhoB consensus motif deduced by the motif-finding program MEME after comparing 98 relevant peaks regions enriched in Δ*pstS*. This analysis reveals a complex 18 nucleotides (nt) PhoB consensus motif, consisting of two pho box repeats (5'-YGTCAYR-3') separated by a conserved A-rich 4 nt spacer (E-value 2.8e-58).

**Supplemental Figure 8. Both phosphate starvation and (p)ppGpp alarmone signaling are required to reinstall motility and the cell cycle-regulated switch in capsulation in cells lacking GcrA.**

(**A**) Motility assays on swarm (0.3%) agar of *WT* and Δ*gcrA::*Ω Δ*gcrB hprK** *pstS** expressing from *pstS* or $pstS^{G61S}$ *in trans* from the vanillate-inducible $P_{van}$ promoter on plasmid pMT335. (**B**) Motility assays on swarm (0.3%) agar of Δ*gcrA::*Ω Δ*pstS*, Δ*gcrA::*Ω Δ*phoD* and Δ*gcrA::*Ω Δ*phoB* cells carrying or not *relA'* under control of the xylose-inducible $P_{xylX}$- promoter at the *xylX* locus. Xylose (0.3%) or/and glucose (0.2%, to repress $P_{xylX}$) were added as indicated. (**C**) Buoyancy of *WT*, Δ*gcrA::*Ω, Δ*phoB*, Δ*spoT*, Δ*phoB* Δ*spoT*, Δ*gcrA::*Ω Δ*phoB*, Δ*gcrA::*Ω Δ*phoB* Δ*spoT*, Δ*gcrA::*Ω Δ*phoB* Δ*spoT* $rpoB^{*H559R}$ and Δ*gcrA::*Ω Δ*phoB* Δ*spoT* $rpoB^{*T561A}$

cells. (**D**) Immunoblots showing steady-state levels of PodJ and FtsN in *WT*, Δ*gcrA::*Ω, *rpoB*$^{H559P}$ and Δ*gcrA::*Ω *rpoB*$^{H559P}$ derivative strains during exponential growth in PYE. MreB serves as a loading control. (**E**) β-galactosidase activities of $P_{mipZ}$-, $P_{ftsN}$-, $P_{pstC}$- and $P_{1149}$-*lacZ* promoter probe plasmids in *WT*, *rpoB*$^{H559P}$, Δ*pstP* Δ*spoT* and Δ*pstP* Δ*spoT* *rpoB*$^{H559P}$ cells measured during exponential growth in PYE. β-galactosidase activities are expressed as percentage relative to *WT* measured in Miller units. Data are from four independent experiments; error bars are standard deviation.

**Supplemental Figure 9. Both phosphate starvation and (p)ppGpp alarmone are required to restore G1 cells population to Δ*gcrA* cells.**

DNA content analysis as determined by FACS (FL1-A channel) of *WT*, Δ*phoB*, Δ*gcrA::*Ω, Δ*phoB* Δ*gcrA::*Ω carrying or not *relA'* under the xylose-inducible $P_{xylX}$- promoter at the *xylX* locus were measured during exponential growth in PYE supplemented with 0.3% of xylose or 0.2% of glucose (to repress $P_{xylX}$) as indicated. The percentages of cells containing >2 chromosomes are also indicated.

**Supplemental Figure 10. *rpoD* is part of the GcrA regulon.**

β-galactosidase activities of $P_{rpoD}$-*lacZ* promoter probe plasmids in Δ*gcrA::*Ω *xylX::*$P_{xylX}$-*gcrA* cell were measured during exponential growth in PYE supplemented with 0.3% of xylose or 0.2% of glucose (to repress $P_{xylX}$) as indicated. β-galactosidase activities are expressed as percentage relative to *WT* measured in Miller units. Data are from four independent experiments; error bars are standard deviation.

**Table S1. *ccrM* Tn-Seq analysis.**

**Table S2. PhoB ChIP-Seq analysis.**

### Supplemental Figure 1

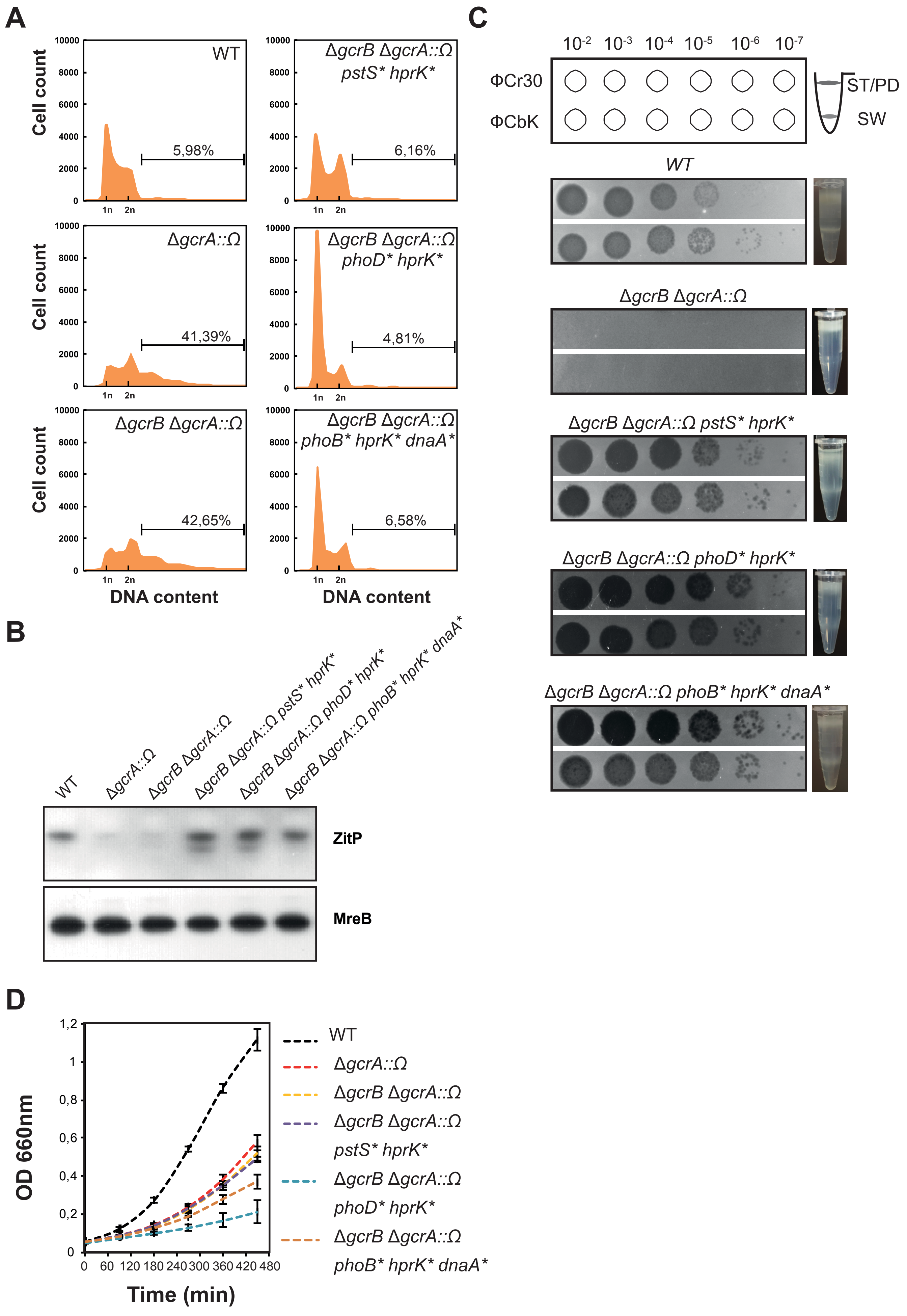

#### Supplemental Figure 2

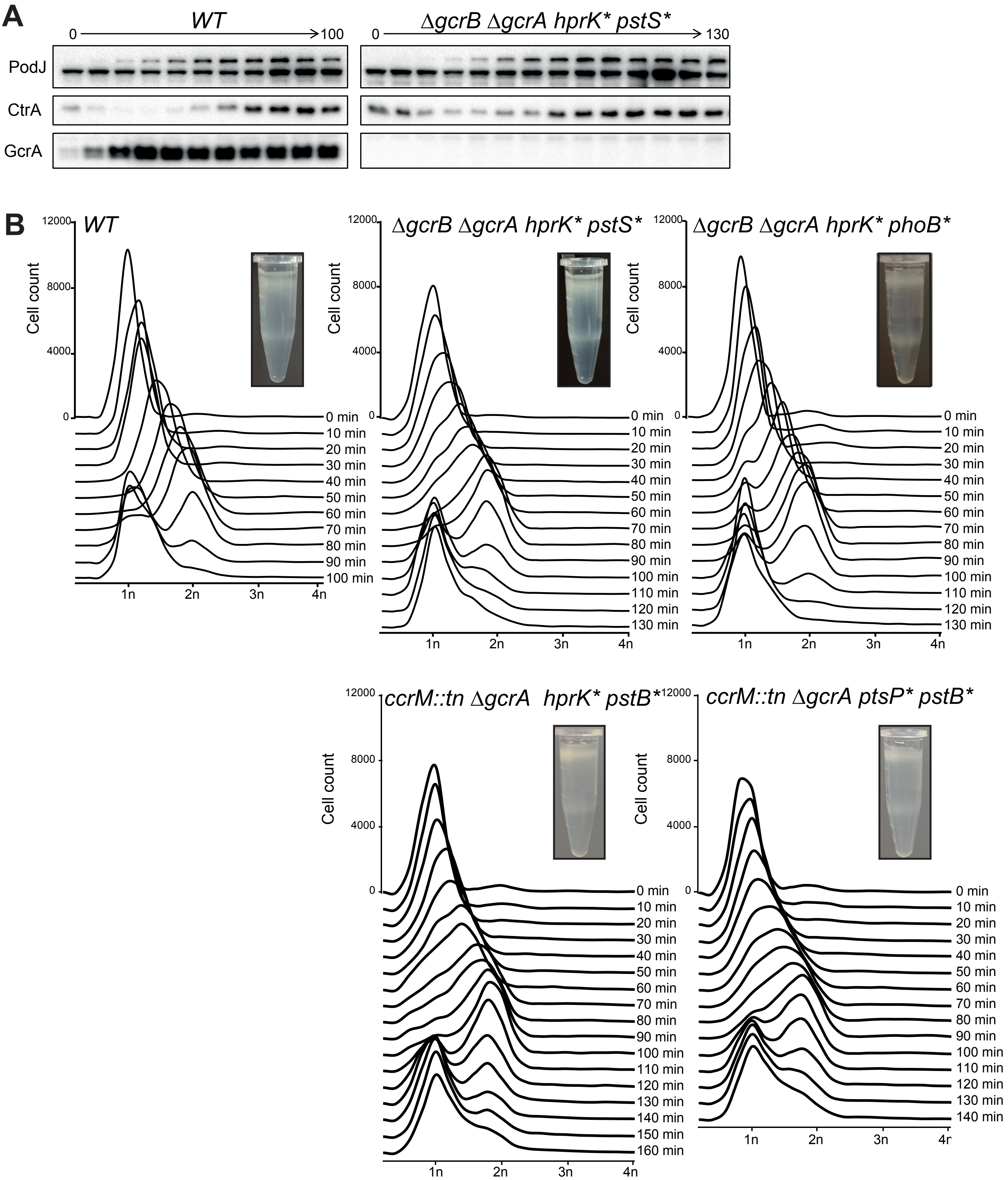

### Supplemental Figure S3

**A**

CDS Tn insertions enrichment in *ΔccrM::Ω*

1000
100
10
0

| Ranking | | |
|---|---|---|
| 1 | *CCNA_01098* | LSU ribosomal protein L35P |
| 2 | *CCNA_01935* | FeS assembly SUF system protein |
| 5 | *CCNA_00321* | LSU ribosomal protein L21P |
| 7 | *CCNA_03971* | LSU ribosomal protein (*ftsZ* promoter region) |
| 9 | *CCNA_01739* | LSU ribosomal protein L9P |
| 14 | *spoT* | Rsh ppGpp hydrolase-synthetase |
| 15 | *sufB* | ABC transporter associated protein SufB |
| 21 | *ccrM* | Adenine specific methyltransferase CcrM |
| 22 | *hprK* | serine kinase/phosphatase HprK |
| 23 | *sufC* | ATP-dependent transporter SufC |
| 28 | *CCNA_01941* | cysteine desulfhydrase/selenocysteine lyase |
| 29 | *gcrA* | cell cycle regualtory protein GcrA |
| 31 | *sufD* | cysteine desulfurase/selenocysteine lyase |
| 46 | *CCNA_01740* | SSU ribosomal protein S18P |
| 55 | *CCNA_01939* | ADP-ribosylglycohydrolase |
| 59 | *CCNA_00240* | PTS system, IIA component |
| 64 | *CCNA_03303* | translation elongation factor TU (EF-Tu) |
| 67 | *sigU* | RNA polymerase ECF-type sigma factor sigU |

**B**

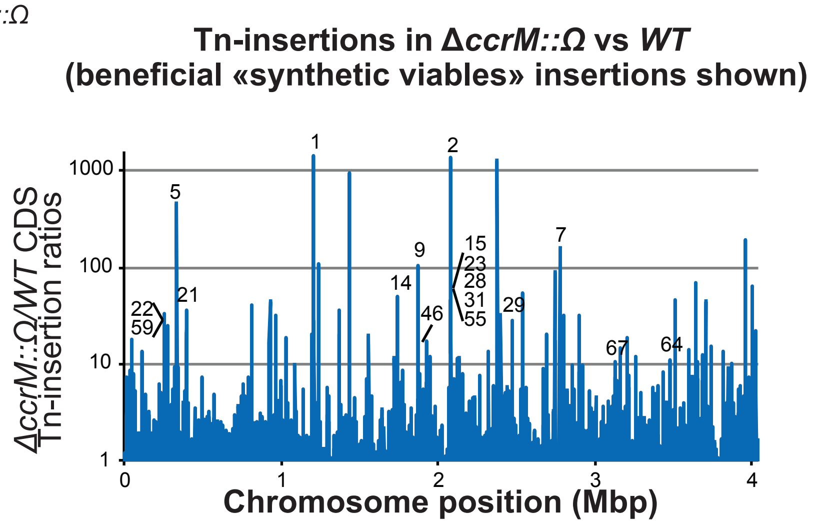


**C**

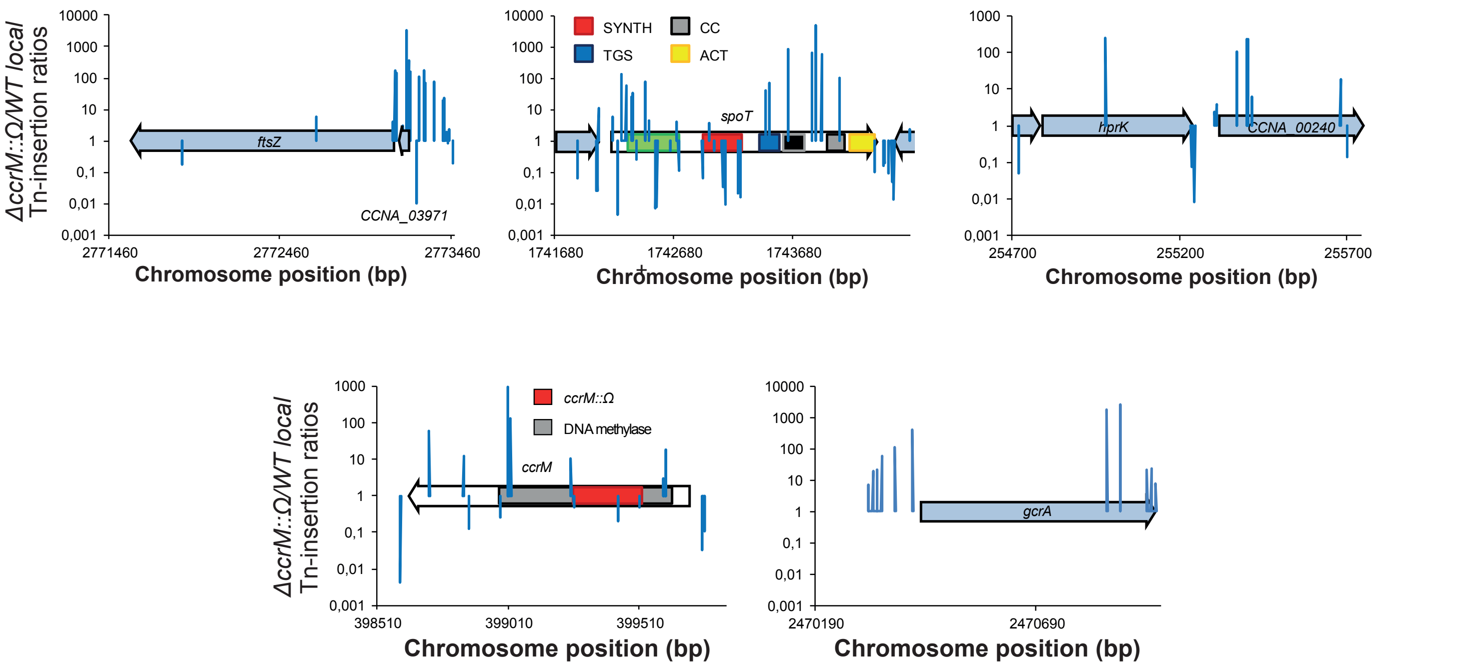

#### Supplemental Figure S4

**A**

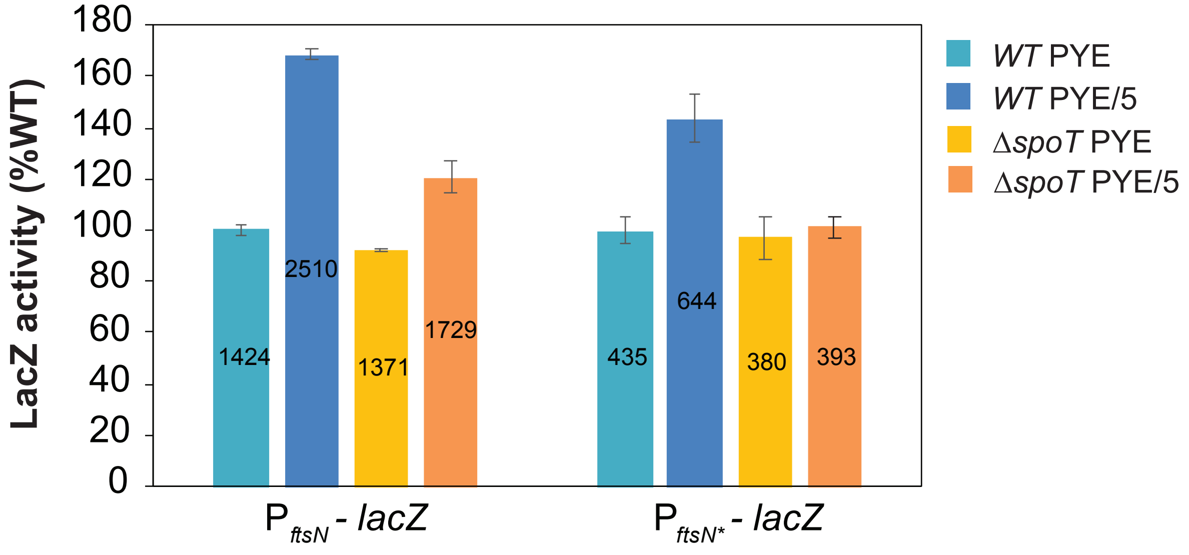


**B**

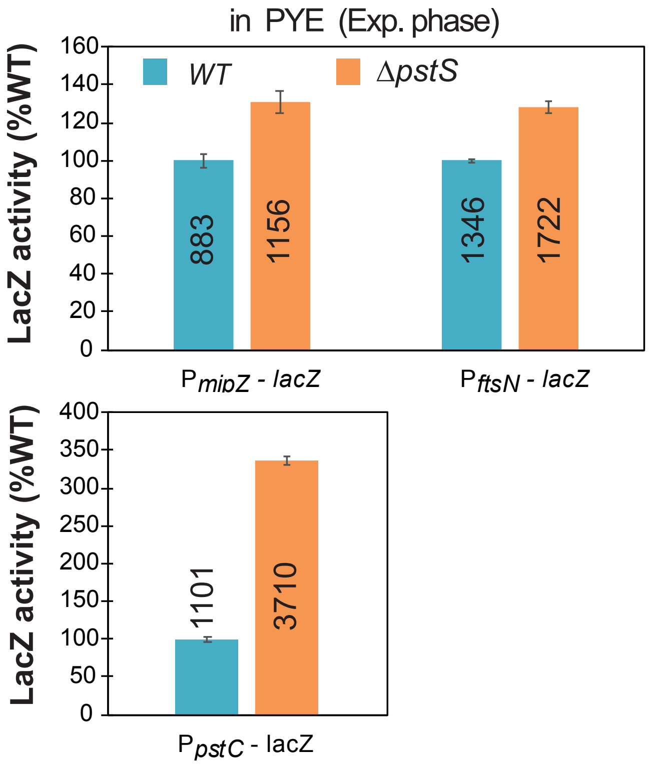

### Supplemental Figure S5

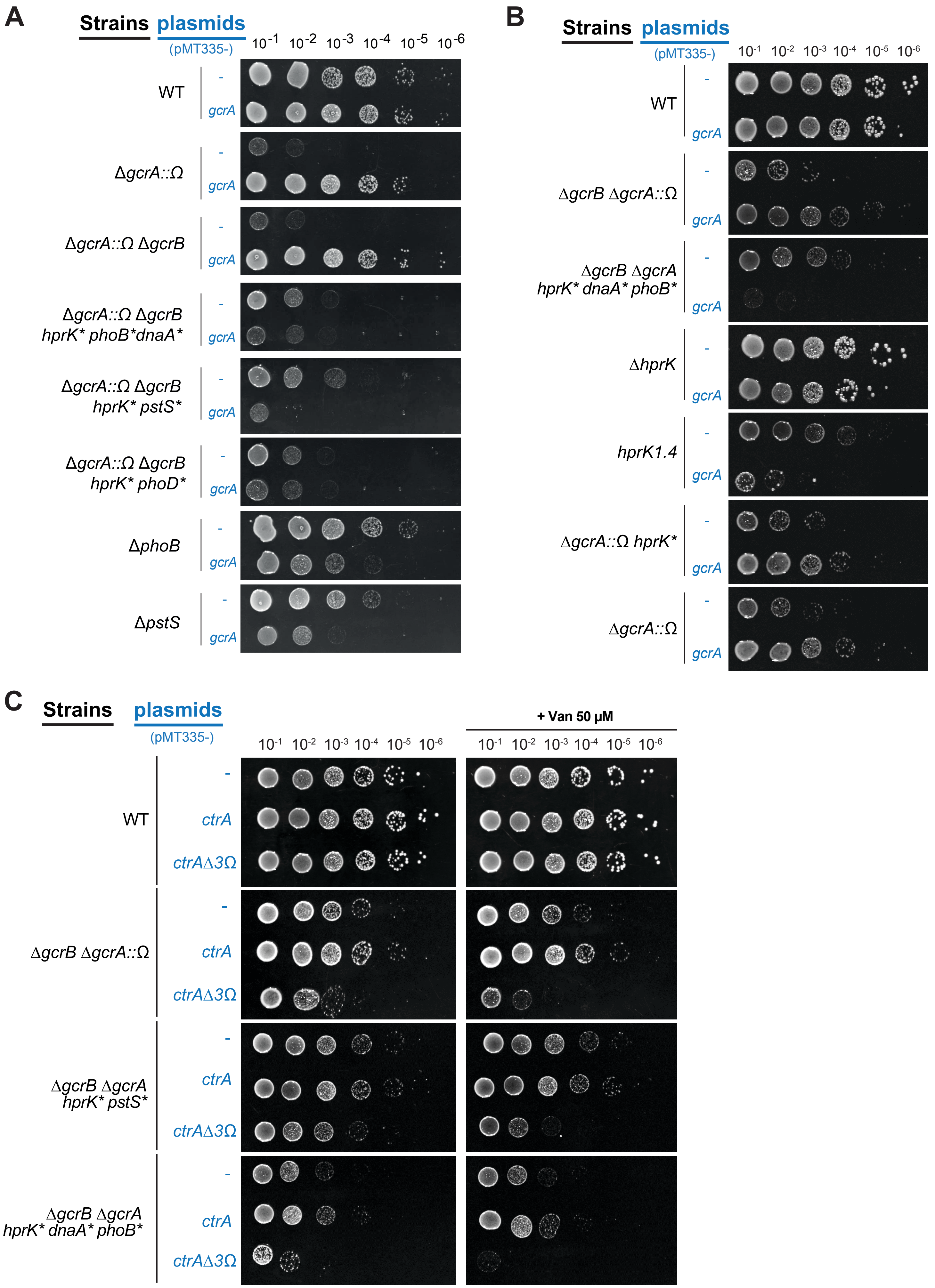

### Supplemental Figure S6

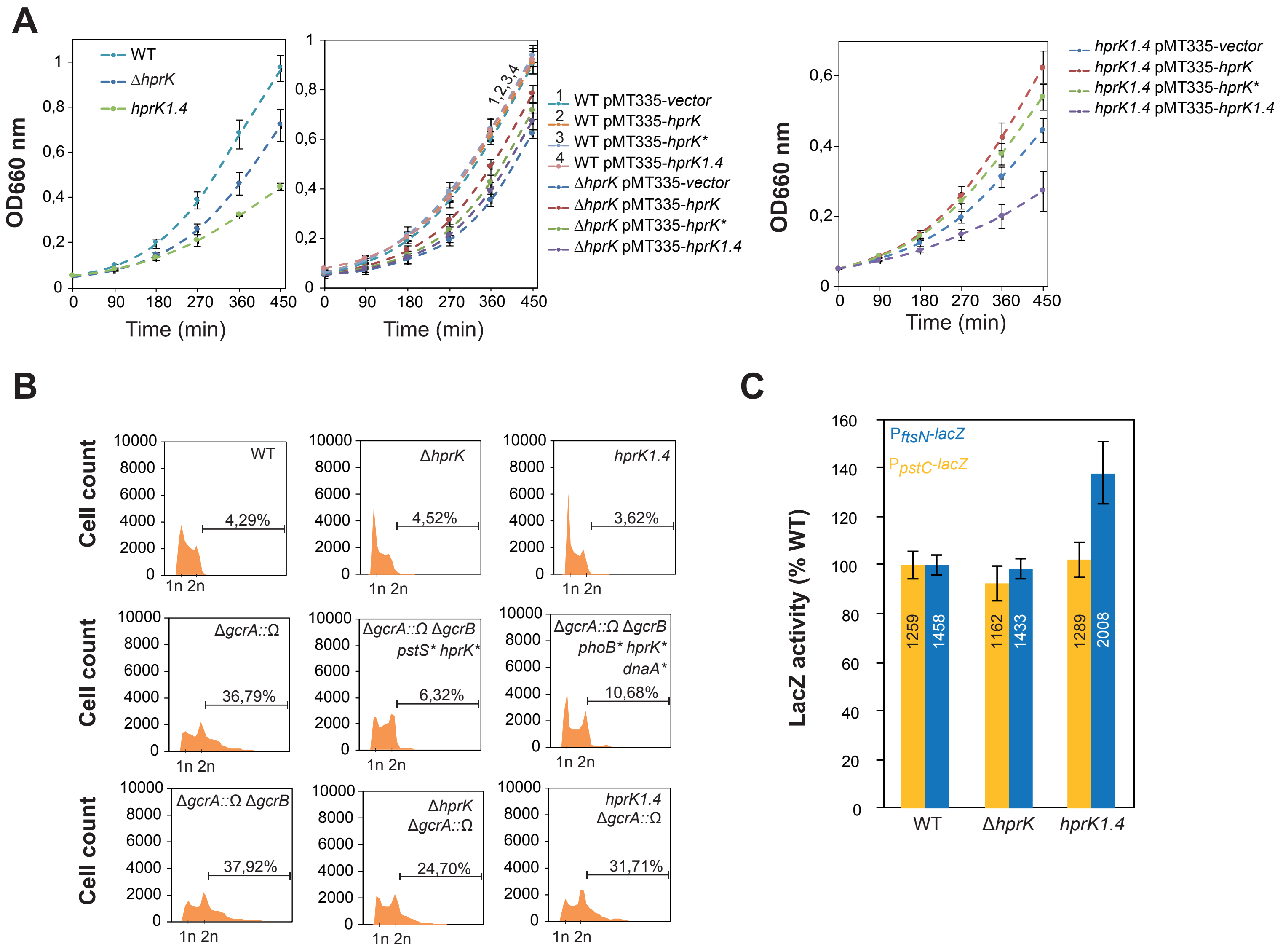

### Supplemental Figure S7

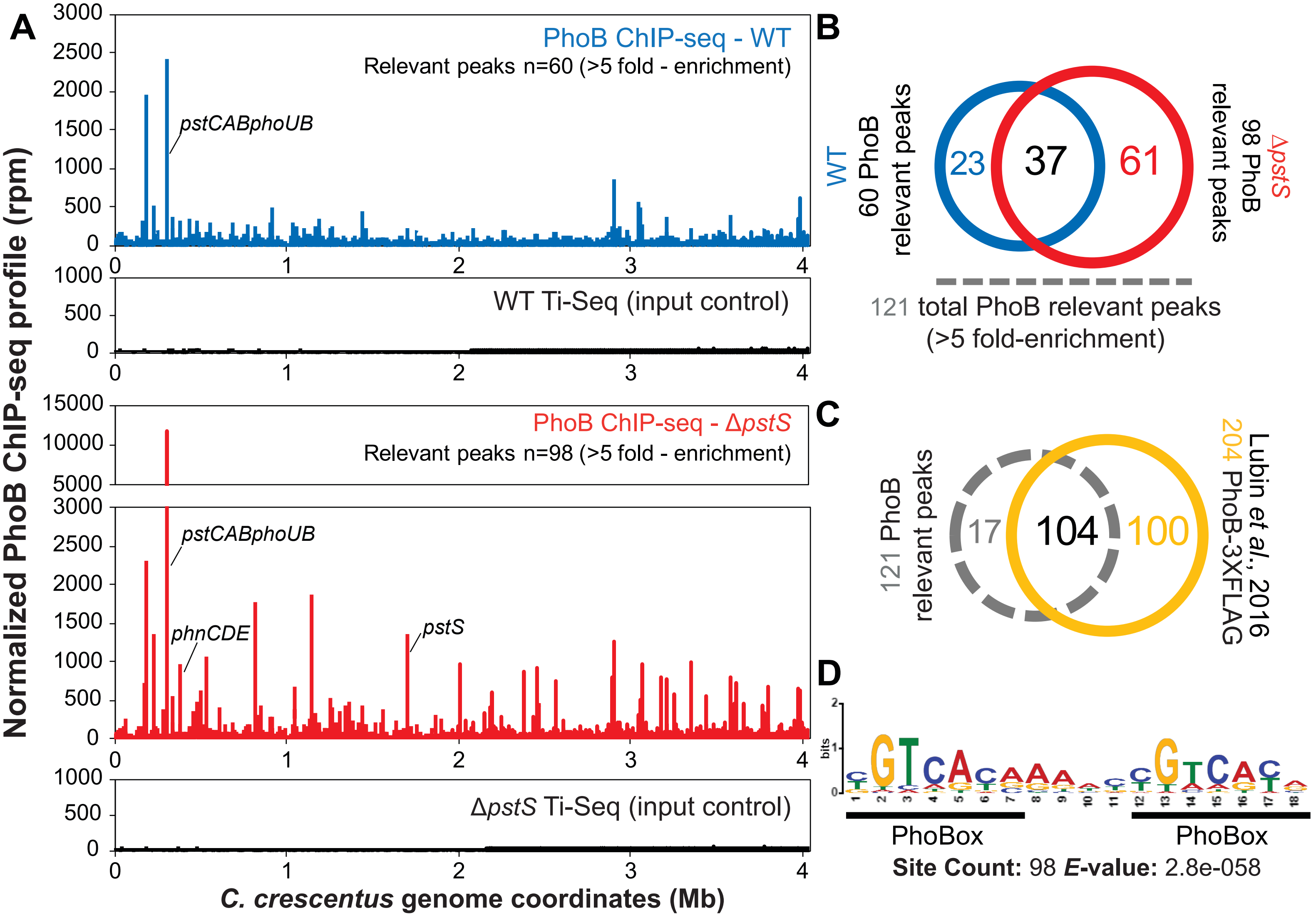

### Supplemental Figure S8

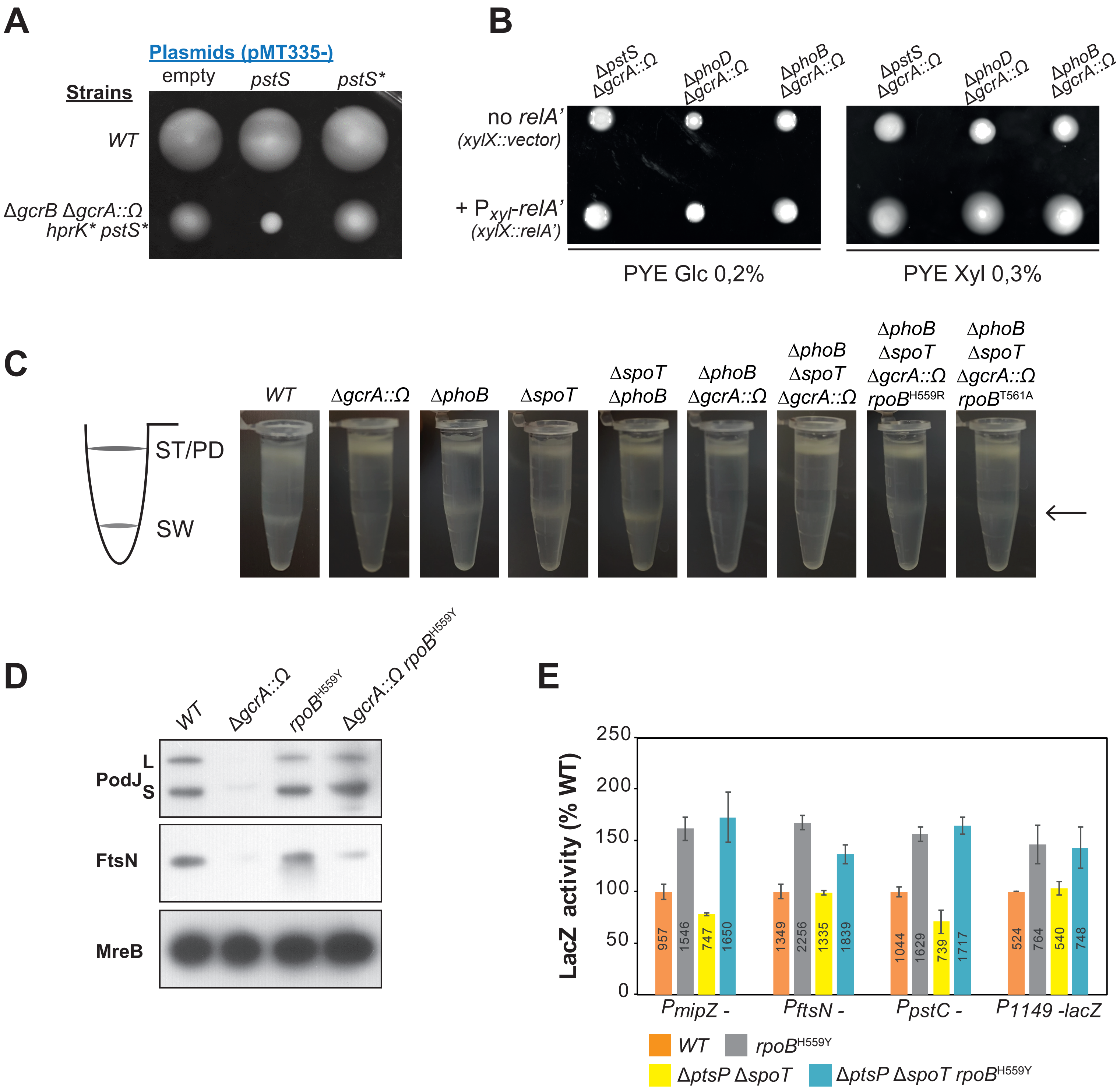

### Supplemental Figure S9

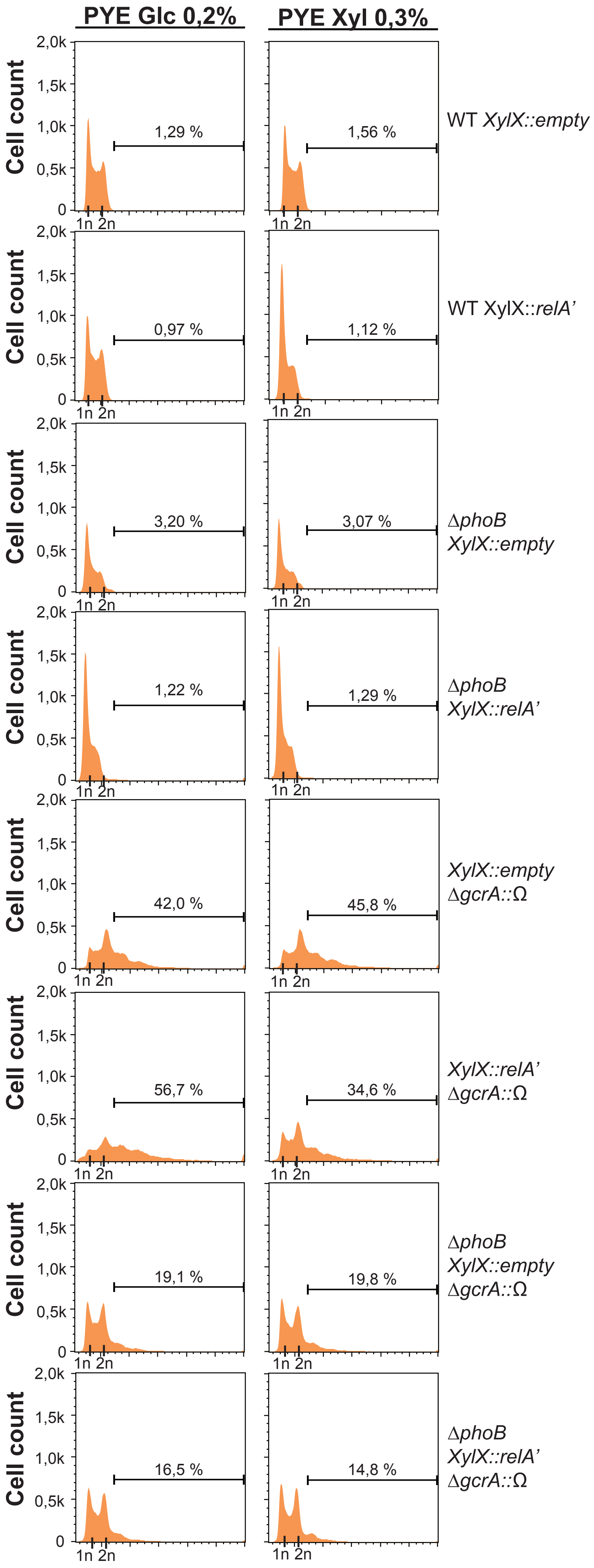

#### Supplemental Figure S10

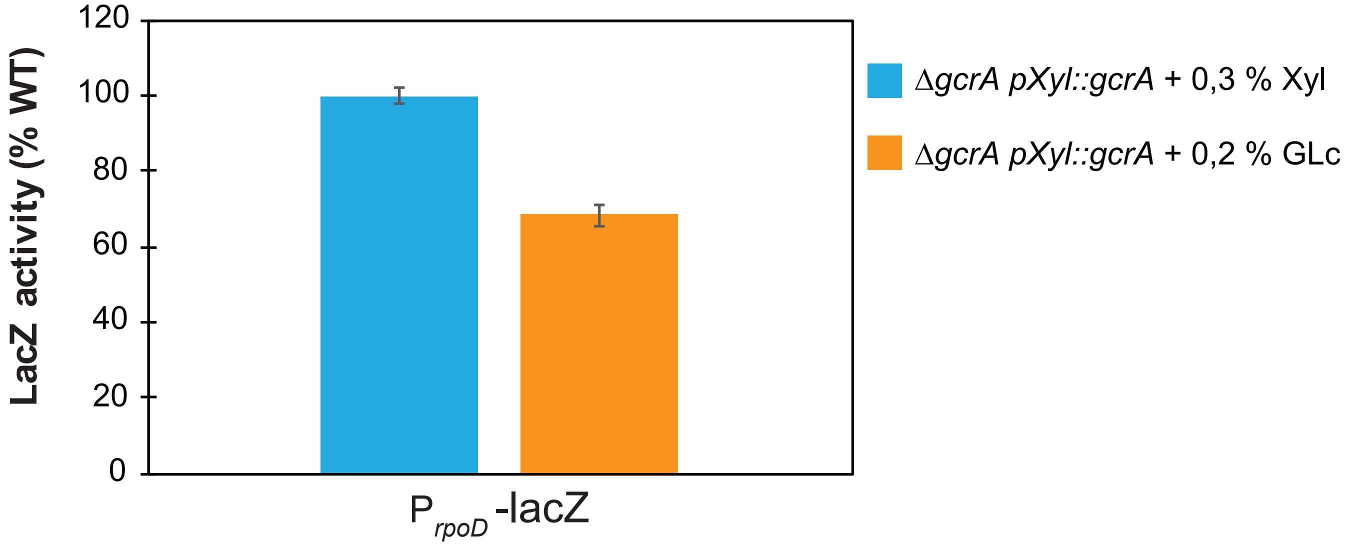
